# Genome-Scale CRISPR Screening reveals novel factors regulating Wnt-dependent regeneration of mouse gastric organoids

**DOI:** 10.1101/2020.07.17.208496

**Authors:** Kazuhiro Murakami, Yumi Terakado, Kikue Saito, Yoshie Jomen, Haruna Takeda, Masanobu Oshima, Nick Barker

## Abstract

An ability to safely harness the powerful regenerative potential of adult stem cells for clinical applications is critically dependent on a comprehensive understanding of the underlying mechanisms regulating their activity. Epithelial organoid cultures accurately recapitulate many features of *in vivo* stem cell-driven epithelial regeneration, providing an excellent *ex vivo* platform for interrogation of key regulatory mechanisms. Here, we employed a Genome-Scale CRISPR Knock-Out (GeCKO) screening assay using mouse gastric epithelial organoids to identify novel modulators of Wnt-driven stem cell-dependent epithelial regeneration in the gastric mucosa. In addition to known Wnt pathway regulators such as *Apc*, we found that knock-out (KO) of *Alk*, *Bclaf3* or *Prkra* supports the Wnt independent self-renewal of gastric epithelial cells *ex vivo*. In adult mice, expression of these factors is predominantly restricted to non *Lgr5*-expressing stem cell zones above the gland base, implicating a critical role for these factors in suppressing Wnt-dependent self-renewal of gastric epithelia. Furthermore, using comprehensive RNA-sequencing analysis, we found that these factors influence epithelial regeneration by regulating Wnt signalling, apoptosis and expression of Reg family genes which could contribute to the epithelial regeneration through JAK/STAT3 pathway.

## INTRODUCTION

The mouse stomach comprises a proximal, non-glandular region not present in humans, and a more distal glandular region that can be anatomically and functionally subdivided into the pylorus and corpus regions (1). The epithelial lining of these glandular regions is composed of multiple tubular invaginations called gastric units, which each comprise a pit, isthmus and base domain and regularly self-renew throughout life. In the pylorus, active stem cell populations at the gland base marked by the Wnt target gene *Lgr5* effect the homeostatic renewal of specialized epithelial cells responsible for secreting protective mucins and hormones such as gastrin that regulate acid and zymogen secretion in the corpus (2). Epithelial renewal in the corpus region is less well understood, but is thought to be largely driven by isthmus-resident stem cell populations (3). Quiescent *Lgr5*-expressing populations at the gland base are rapidly recruited to function as reserve stem cells following the injury-driven loss of the homeostatic stem cell pool (4). Lgr5 is instrumental in modulating R-spondin mediated regulation of canonical Wnt signalling on stem cells in the glandular stomach as an essential prerequisite to balance self-renewal and differentiation *in vivo* (5).

Development of 3D epithelial culture systems capable of supporting the growth of highly stable, self-renewing primary tissues that more accurately recapitulate the composition and functionality of the tissue of origin has revolutionized the study of epithelial biology (6, 7). In the stomach, epithelial organoid formation is selectively driven by primary glandular *Lgr5*-expressing cells, further underscoring their endogenous stem cell identity (3, 4). Like their *in vivo* counterparts, the *ex vivo* stem cells display an absolute dependence on canonical Wnt signals to regulate their ability to orchestrate self-renewal and differentiation within the organoids. However, despite these observations, detailed mechanistic insight into the regulation of stem cell renewal versus differentiation by Wnt signalling in the stomach is currently lacking.

Whilst the organoid culture system doesn’t fully mimic the homeostatic *in vivo* microenvironment, it is by far the most physiological *ex vivo* model of stem cell-driven epithelial regeneration available (8). Together with the ready accessibility of its genome for performing phenotypic screening, the organoid provides a powerful model for dissecting the mechanism of Wnt-driven regulation of epithelial regeneration in the stomach.

Genome-wide, targeted loss-of-function pooled screens using the clustered, regularly interspaced, short palindromic repeats (CRISPR)-associated nuclease Cas9 (CRISPR/Cas9) in human and mouse cells provide an alternative screening system to RNA interference (RNAi) (8). The Genome-scale CRISPR KO pooled library was firstly reported by Sanjana N.E. *et al*. (9), and several libraries subsequently developed (11, 12, 13). These genome-wide screening techniques have been applied to several cancer cell lines to clarify the mechanism which induces malignancy (14, 15). However, an application for the normal gastric organoid to identify the mechanism which regulates stem cell-driven epithelial regeneration has not been reported so far.

Here, we combined the GeCKO screening system and gastric organoid culture techniques to investigate the underlying mechanism by which Wnt signalling regulates self-renewal versus differentiation to maintain the integrity of the gastric epithelium. We established corpus and pyloric organoids from mouse normal stomach and confirmed that the Wnt signal was necessary to maintain the self-renewal of those organoids. We subsequently applied the GeCKO screening method to those normal gastric organoids. Using next-generation sequencing, we found that *Alk*, *Bclaf3* and *Prkra* are novel Wnt signal suppressors present on non-stem cell compartments. We also found that these factors regulate the differentiation of gastric epithelium by suppressing Wnt pathway, CD44 and *Reg* family genes, which influence epithelial cell regeneration via modulation of JAK/STAT3 signalling.

## MATERIAL AND METHODS

### Mice

Lgr5-DTR-EGFP mice have been described previously (16). The line is bred as heterozygotes. All animal experimental protocols were approved by the Committee on Animal Experimentation of Kanazawa University.

### Preparation of L-WRN conditioned medium

L-WRN cell line was purchased from ATCC (ATCC CRL-3276) and maintained in DMEM (Wako) with 10 % FBS and 1x Penicillin /Streptomycin (Nacalai Tesque) (17). L-WRN conditioned medium was prepared according to the protocol provided by ATCC.

### Organoid culture

Organoid culture was performed as described previously with minor modifications (4). Briefly, 2-3 mm tissue fragments from murine corpus and pylorus were incubated in 1xPBS supplemented with 5 mM and 10 mM EDTA (Corning) at 4 °C for 2 and 3 hours respectively. Glands were isolated by repeated pipetting of finely chopped tissues in ice-cold 1xPBS supplemented with EDTA. 1xPBS containing isolated glands was filtered through 100 μm filter mesh and centrifuged at 720g at 4 °C for 3 min. Whole gastric glands were embedded in GFR Matrigel (Corning) and cultured in basal media (Advanced DMEM/F-12 media with 10 mM HEPES, 2mM Glutamax, 1X N2, 1X B27 (Invitrogen), N-acetyl-cysteine (Sigma-Aldrich) supplemented with growth factors, EGF (Peprotech), Gastrin (Wako), FGF10 (Peprotech), L-WRN conditioned medium and passaged every 3 days. For passage, organoids were collected with Recovery Solution (Corning), dissociated by gentle pipetting and transferred to Matrigel. For low Wnt medium, the L-WRN condition medium was reduced to 5 % and supplemented with Noggin (Peprotech) and FBS.

For organoid out-growth assays, organoids were cultured for 3 passages in 5 % L-WRN supplemented with Noggin. Parental organoids were cultured in 50 % L-WRN conditions. Lgr5^+^ cells were ablated by treatment with 500 nM DT after 3 days, before being dissociated. 5,000 cells were seeded in a well (3 technical triplicates × 2 biological replicates). DT treatment was continued throughout the assay.

Canonical Wnt signal inhibitors, IWR-1-endo (Wako) and XAV939 (Sigma-Aldrich) were added to the medium at 5 μM and 1 μM respectively. The porcupine inhibitor, IWP-2 (Wako) was added to the medium at 2 μM. Recombinant Human REG1A, REG3b and REG3G (R&D) were added to 500 μg/ml.

### HEK293T cell culture

HEK293T cell line was maintained in DMEM (Wako) with 10 % Fetal Bovine Serum (FBS) (Sigma-Aldrich) and 1x Penicillin /Streptomycin (Nacalai Tesque). Passaging was performed using TrypLE (ThermoFisher Scientific) according to standard protocols.

### Preparation of lenti-GeCKO library

Mouse GeCKOv2 CRISPR knockout pooled library was a gift from Feng Zhang (Addgene # 1000000052). To prepare lentivirus particles, the Mouse GeCKOv2 CRISPR knockout pooled library plasmids were co-transfected with packaging plasmids pCMV-VSV-G and pCMV-dR8.2 dvpr (Gifts from Bob Weinberg, Addgene plasmid #8485 and #8455) into HEK293T cells. Briefly, 80 % confluent HEK293T cells in 100 mm dishes were transfected in OptiMEM (Life Technologies) using 6 μg of Mouse GeCKOv2 CRISPR knockout pooled library plasmids, 1.5 μg pCMV-VSV-G, 4.5 μg pCMV-dR8.2 dvpr, 60 μl of 1 ug/ml PEI MAX (Polysciences Inc.). After 16 hours, media was changed to Advanced DMEM/F-12 with 1x B-27 and 1x GlutaMax. After 72 hours, viral supernatants were harvested and centrifuged at 1,000 rpm at 4 °C for 5 min to pellet cell debris. The supernatant was filtered through a 0.45 μm membrane filter (Sartorius) and aliquots were stored at −80 °C until use.

The GeCKO library is divided into two sub-libraries, A and B. Both libraries comprises three sgRNAs targeting each gene, as well as 1,000 non-targeting control sgRNAs. Library A also contains sgRNAs targeting non-coding micro RNAs. Normal corpus and pylorus organoids were infected at a low MOI (<0.4) to ensure that most cells receive only 1 viral construct with high probability (18). Briefly, organoids were dissociated by TrypLE (ThermoFisher Scientific) and cells were mixed with the lentivirus solution and 8 μg/ml polybrene (Sigma Aldrich). The sample was centrifuged at 600 g at 32 °C for 60 mins and incubated at 37 °C for 6 hours. After incubation, cells were collected and embedded in matrigel with medium containing Y-27632 (Wako) and CHIR99021 (Cayman Chemical).

To determine MOI, cells were counted and each well was split into duplicate wells. One replicate received 1 μg/mL puromycin (Invivogen). After 3 days, cells were counted to calculate percent transduction. Percent transduction is calculated as cell count from the replicate with puromycin divided by cell count from the replicate without puromycin multiplied by 100. The virus volume yielding a MOI closest to 0.4 was chosen for large-scale screening.

For the large-scale screening, 24 hours after transduction, organoids were selected with 1 μg/ml puromycin and 5 % L-WRN supplemented media for 3 passages. After 3 passages, only organoids transduced with lentiCRISPR constructs and can proliferate in the absence of Wnt signal were preserved. These organoids were collected for use in subsequent analyses. We independently repeated this screening steps 3 times.

### Identification of integrated gRNA

#### Amplicon sequencing

The primer sets HT-gLibrary-Miseq_150bp-PE-F: TCGTC GGCAGCGTCAGATGTGTATAAGAGACAGcttgaaagtatttcgatttcttgg and HT-gLibrary-Miseq_150bp-PE-R: GTCTCGTGGGCTCGGAGATGTGTATAAGAGACAGactcggtgccactttttcaa were used to amplify the gRNA locus. The PCR condition was 95 °C (20 sec), 55 °C (20 sec), and 72 °C (30 sec) for 26 cycles. Q5 Taq polymerase (New England Biolabs) was used. Secondary PCR was performed using Nextra XT index primers (Illumina). The secondary PCR condition was 95 °C (20 sec), 55 °C (20 sec), and 72 °C (30 sec) for 12 cycles. PCR products were purified using a PCR purification kit (Hokkaido System Science) and sequenced on a Next-Seq 500 (Illumina) system (75-bp single-end reads). Sequence reads were trimmed and quality-filtered using Trim Galore! (version 0.4.4), Trimmomatic (ver. 0.36), and Cutadapt (ver. 1.16). Frequency of occurrence of each contig sequence was calculated using the table function of R (version 3.4.1).

#### Manual sequencing

Inserted gRNA sequences were amplified with the following primers: LentiCRISPR v2 gRNA scaffold F; 5’-TGCATGCTCTTCAACCTCAA −3’, LentiCRISPR v2 gRNA scaffold R; 5’-CCAATTCCCACTCCTTTCAA-3’. PCR fragments were gel extracted and cloned with the pGEM-T easy Vector system (Promega) and transfected into XL10Gold competent cells (Agilent). After Blue/White selection, only white colonies which contain an amplified sequence were picked and plasmids subsequently extracted. Cloned fragments were sequenced by the standard sanger sequencing method using T7 and SP6 universal primers. We sequenced at least 100 plasmids from each screening.

#### KO of individual genes using CRISPR/Cas9

Identified gRNAs in organoids which survived under low Wnt conditions were cloned into pX330-U6-Chimeric_BB-CBh-hSpCas9 (a gift from Feng Zhang, Addgene #42230) and transfected into organoids and HEK293T cells with Lipofectamine LTX or Lipofectamine 2000 (ThermoFisher Scientific) respectively according to manufacturer instructions. Plasmids were co-transfected with a CAG-EGFP-IRES-Puro^r^ vector and cells were selected by 1 μg/ml puromycin for 3 days. Additional gRNAs which were not listed in the GeCKO library were designed by the CHOPCHOP software. KO efficiency was confirmed by Western blotting. gRNAs used this study are listed in Supplementary Table 1.

#### Overexpression of *Prkra*, *Eif2ak2*, *Reg1*, *Reg3β*, *Reg3γ*

*Prkra* and *Eif2ak2* genes were amplified from a cDNA pool prepared from mouse normal gastric organoids. Reg1, Reg3β and Reg3γ cDNA plasmids were provided by the RIKEN BRC through the National BioResource Project of the MEXT/AMED, Japan. cDNAs were cloned into pPBCAG-IRES-Puro^r^, a PiggyBac-based puromycin resistant constitutive expression vector or pPBhCMV*1-pA, a PiggyBac-based inducible expression vector (Gifts from Dr. H. Niwa). These vectors were transfected into organoids derived from normal stomach of Lgr5-DTR-EGFP mice, together with pPyCAG-PBase vector and pPBCAG-rtTA3G-IN vector as described previously (4). After 1 week of Puromycin (1 μg/ml; InVivogen) or G418 (500 μg/ml; Wako) selection, pooled clones were used for subsequent experiments.

### Histology

#### Immunohistochemistry (IHC) and immunofluorescence (IF)

IHC was performed according to standard protocols. Briefly, tissues were fixed in 4 % paraformaldehyde/PBS (w/v) overnight and processed into paraffin blocks. 4 μm sections from the paraffin blocks were deparaffinated and rehydrated, followed by antigen retrieval via heating in a microwave in standard 10mM citric acid pH6 buffer. Primary antibodies used were anti-Reg1α (1:100, Abcam), anti-Reg3γ (1:100, LSBio). To detect signals, ImmPRESS Reagent (Vector laboratories) was used. IHC sections were dehydrated, cleared, and mounted with EUKITT mounting medium (ORSAtec). Immunostaining was performed on a minimum of three biological replicates and representative images of the replicates were included in the manuscript. Sections were imaged on a microscope (Leica)

For organoid staining, IF was performed in Matrigel. Briefly, organoids were fixed in 4 % paraformaldehyde/PBS (w/v) for 20 min and permeabilized in 0.5 % Triton-x100/PBS at 4 °C for 1 hour. After blocking with 10 % normal donkey serum, Primary antibodies used were anti-Cleaved Caspase-3 (1:200, Cell Signalling), anti-Ki67 (1:100, BioScience), anti-CD44 (1:200, Millipore). To detect signals, an Alexa 594 conjugated second antibody (1:500; Invitrogen) was used. Stained organoids were mounted with Vectashield antifade containing DAPI (DAKO) for imaging. Immunostainings were performed on a minimum of three biological replicates and representative images of the replicates were included in the manuscript. Sections were imaged on a confocal microscope (Leica).

#### Hematoxylin and eosin (H&E)

H&E staining was performed on FFPE sections according to standard protocols.

#### RNA *In Situ* hybridization (ISH)

RNA ISH was performed using RNAscope 2.5 High Definition Brown Assay (Advanced Cell Diagnostics) according to the manufacturer’s instructions. DapB was used as a negative control for all the RNAscope experiments. *In situ* hybridization and imaging was performed on a minimum of three biological replicates and representative images of the replicates were included in the manuscript. Probes for each gene were obtained from Advanced Cell Diagnostics Inc.

#### RNA-sequencing

RNA sequence library preparation, sequencing, mapping, gene expression and gene ontology (GO) enrichment analysis were performed by DNAFORM (Yokohama, Kanagawa, Japan). Quality of total RNA was assessed by Bioanalyzer (Agilent) to ensure that RIN (RNA integrity number) was above 7.0. Double-stranded cDNA libraries (RNA-seq libraries) were prepared using SMARTer® Stranded Total RNA-Seq Kit v2 – Pico Input Mammalian (Clontech) according to the manufacturer’s instruction. RNA-seq libraries were sequenced using a HiSeq 2000 (Illumina), as 150bpPE. Obtained reads were mapped to the mouse mm9 genome using STAR (version 2.7.2b). Reads on annotated genes were counted using featureCounts (version 1.6.1). FPKM values were calculated from mapped reads by normalizing to total counts and transcript. Differentially expressed genes were detected using the DESeq2 package (version 1.20.0). The list of differentially expressed genes detected by DESeq2 (basemean > 5 and fold-change < 0.25, or basemean > 5 and fold-change > 4) were used for GO enrichment analysis by clusterProfiler package (19).

#### β-Catenin Knock-down

ON-TARGET plus SMARtpool siRNAs for β-catenin and ON-TARGET plus SMARTpool non-targeting siRNAs pool (Horizon Discovery) were transfected into organoids at 1 μM. Briefly, organoids were dissociated by TrypLE (ThermoFisher Scientific) and cells were mixed with siRNA cocktails and Lipofectamine RNAi Max (ThermoFisher Scientific). The cells were centrifuged at 600 g at 32 °C for 60 mins and incubated at 37 °C for 6 hours. After incubation, cells were collected and embedded in matrigel with medium containing Y-27632 (Wako) and CHIR99021 (Cayman Chemical). After 24 hours, knock-down (KD) efficiency was confirmed by qPCR. To evaluate cell growth, organoids were maintained for 3 passages without Y-27632 and CHIR99021.

#### TOPFlash assay

M50 Super 8x TOPFlash and M51 Super 8x FOPFlash plasmids (Gifts from Randall Moon, Addgene #12456 and 12457) were transfected into HEK293T cells with Lipofectamine 2000 (ThermoFisher Scientific) according to the manufacturer’s instruction. Next day, luciferase activity was measured using the ONE-Glo luciferase assay system (Promega). Protein quantification was also performed using Pierce 660 nm protein assay reagents (ThermoFisher Scientific). Luciferase activity was normalized against total protein.

#### qPCR

Total RNA was extracted from organoids by RNAeasy Mini Kit or Micro Kit (Qiagen) and cDNA generated with PrimeScript RT reagent Kit (Perfect Real Time) (TAKARA) according to manufacturer’s instructions. qPCR was performed with a minimum of three biological replicates per gene using THUNDERBIRD SYBR qPCR Mix (TOYOBO) according to the manufacturer’s instructions and ran on Mx3000P Real-Time QPCR System (Agilent). Analysis was carried out using double CT method. Primers used this study are listed in Supplementary Table 1.

#### Western blotting

Western Blotting was performed according to standard protocols. Cells were lysed in lysis buffer supplemented with protease inhibitors (Sigma-Aldrich) and a phosphatase inhibitor (ThermoFisher Scientific). Lysates were centrifuged at 12,000 rpm for 5 min at 4 °C. Supernatants were quantified using Pierce 660 nm Protein Assay Reagent (ThermoFisher Scientific). Lysates were denatured at 95 °C for 5 minutes before gel electrophoresis on 8-12 % acrylamide gels. Proteins were then transferred to PVDF membranes at 20 V, 400 mA for 150 min at 4 °C. Membranes were blocked by incubation with 5 % skimmed milk/1x TBS-T. Primary antibodies used: anti-β—Actin (1:2000, Sigma Aldrich), anti-Alk (1:100, SantaCruz), anti-Bclaf3 (1:300, Biorbyt), anti-Prkra (1:1000, GeneTex), anti-Non-Phospho (active) β-catenin (1:1000, Cell Signaling), anti-β-Catenin (1:1000, Cell Signaling), anti-elF2α (1:1000, Cell Signaling), anti-phospho-elF2α (1:1000, Cell Signaling). Membranes were developed by Amersham ECL select (GE lifesciences) and imaged using ImageQuant LAS 4000 (GE lifesciences).

#### FACS analysis

Organoids were dissociated in TrypLE (ThermoFisher Scientific) for 5 min at 37 °C. Digestion was quenched by DMEM 10 % FCS medium. The suspension was centrifuged at 300 g for 5 min. The pellet was resuspended in 1xPBS supplemented with 2 % fetal bovine serum (FBS, Hybclone) and 1 μg/mL 7-AAD (Life Technologies), filtered through a 40 μm strainer, and analysed by BD FACS CantoII or BD FACS AriaIII (BD Biosciences).

#### Statistical methods

Data were quantified and depicted as the mean ± standard deviation (S.D.). Statistical analysis was performed using an unpaired *t*-test. Significant difference is denoted as ∗p < 0.05.

## RESULTS

### Establishment of Genome-scale CRISPR Knock-Out (GeCKO) screening system using gastric organoids

To decipher the underlying mechanisms regulating Wnt-dependent epithelial regeneration in the stomach, we established a Genome-wide CRISPR Knock-Out (GeCKO) screen using normal mouse gastric organoids derived from both the corpus and pylorus regions (**Figure 1a**).

**Figure 1.**
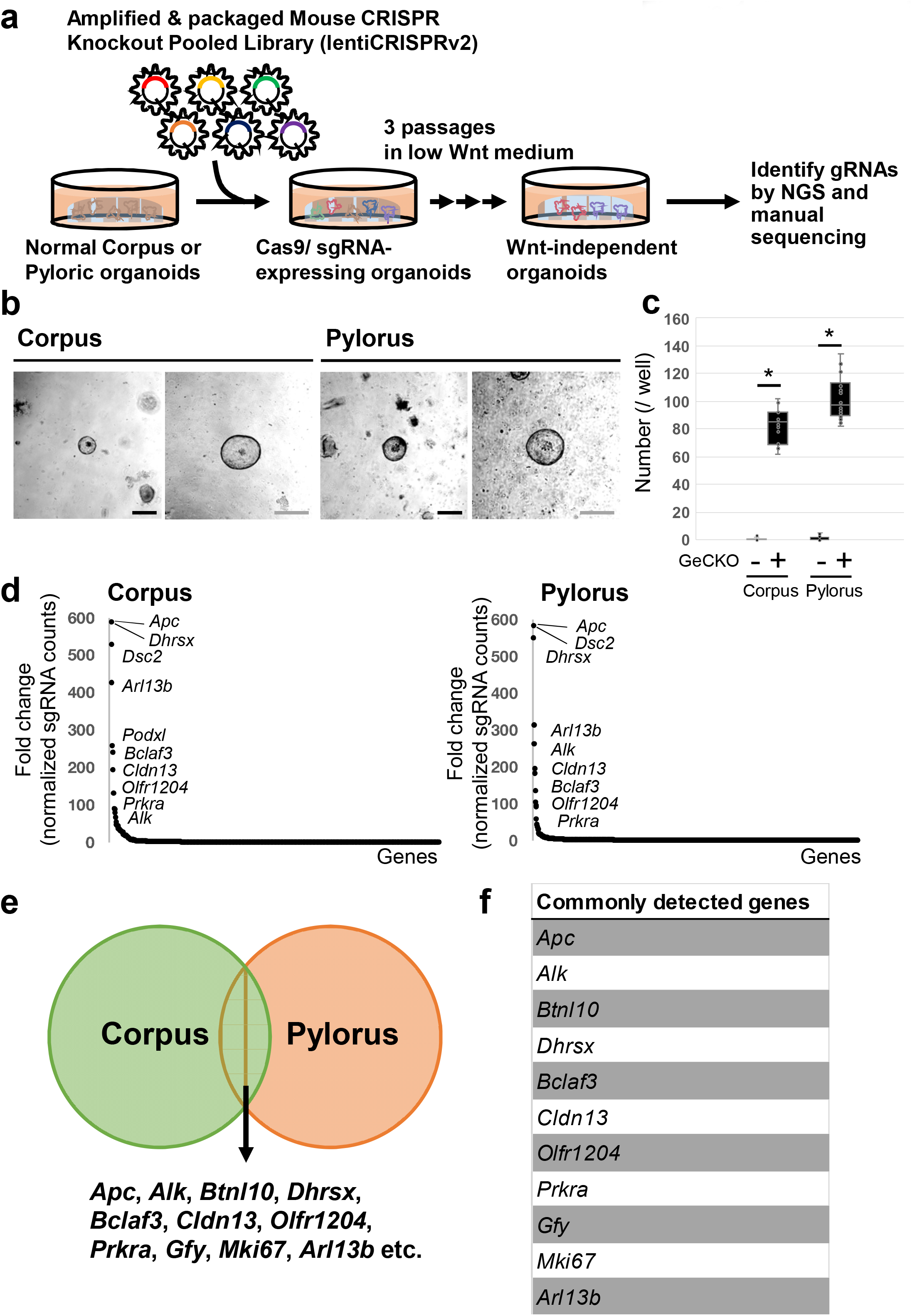
Establishment of Genome-scale CRISPR KO screening in normal gastric organoids. (a) A scheme of the GeCKO screening method to reveal novel regulators of Wnt-dependent epithelial regeneration. (b) Images of representative gastric organoids grown in the low Wnt medium after infection of the GeCKO library. In total, 6 independent experiments were performed. Scale bar = 50 μm. (c) Wnt-independent organoid number per well (N = 6). * p < 0.05. (d) Enriched gRNA targeted genes from results of amplicon-sequencing. (e) Enriched gRNA targeted genes from results of TA cloning followed by sanger sequencing. (f) Commonly listed genes from both amplicon-sequencing and TA cloning-Sanger sequencing.

We first established normal corpus and pyloric gastric organoids from Lgr5DTR-EGFP reporter mice as shown in **Supplementary Figure 1a**. Consistent with previous reports, these organoids could be cultured more than 3 months without major morphological change, confirming their long-term stability (**Supplementary Figure 1b**). Importantly, the organoids could be propagated from single cells, further underscoring the long-term maintenance of epithelial stem cells in this culture system (**Supplementary Figure 1c**).

Next, to confirm which signal is essential for the maintenance of those organoids, we individually depleted the growth factors present in the culture medium. We confirmed that Wnt stimuli are essential for maintaining the growth of normal gastric organoids (**Supplementary Figure 1d**). This is consistent with the *in vivo* situation, where it has been reported that the Wnt/R-spondin pathway is essential for the maintenance of stem cell-driven epithelial renewal (5). To better define the underlying mechanism by which Wnt/R-spondin regulates stem cell-driven epithelial regeneration in the stomach, we decided to perform genome-wide GeCKO screening in gastric organoids under Wnt/R-spondin depleted conditions.

The GeCKO library comprises two libraries, A and B. Each library contains 3 independent sgRNAs for 20,611 individual protein-coding genes and 1,000 control non-target gRNAs. For this study, we used library A as it also contains sgRNAs targeting 1,175 miRNAs and using library B doesn’t increase genome coverage. To perform the genome-wide KO screening, we determined multiplicity of infection (MOI) of lenti-GeCKO libraries for the screening (**Supplementary Figure 2**). According to a previous report (18), we decided to use 50 % lentivirus solution for the large-scale screen.

**Figure 2.**
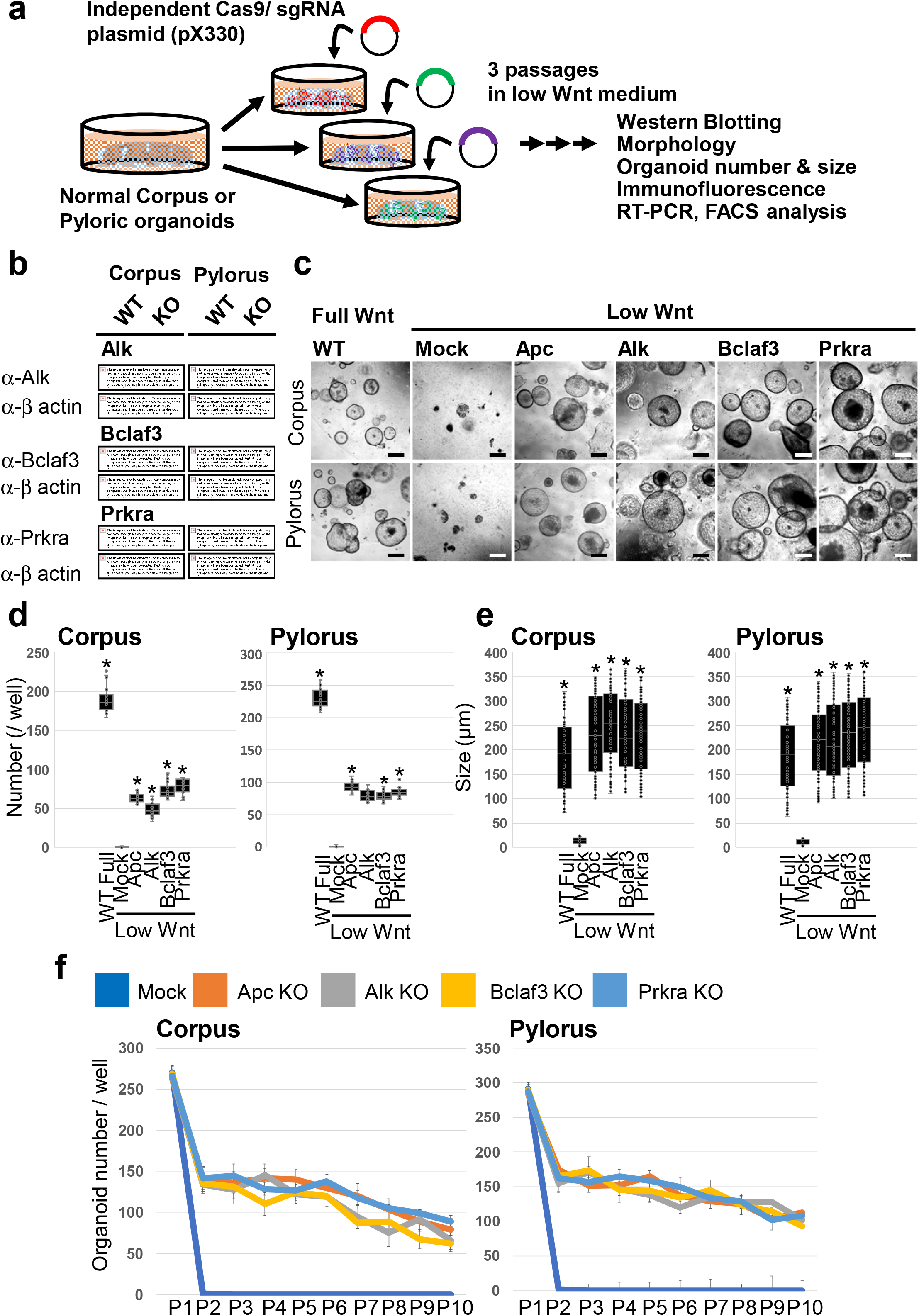
KO of *Alk*, *Bclaf3* and *Prkra* supports organoid survival under low Wnt conditions. (a) Validation of GeCKO screening results. Single sgRNAs from GeCKO library was cloned into the CRISPR/Cas9 plasmid (pX330) and transduced into gastric organoids. Organoids were selected under low Wnt conditions for 3 passages. (b) Confirmation of the KO efficiency of each gene by Western Blotting. (c) Representative images of single-gene KO organoids which proliferated under low Wnt conditions. Scale bar = 200 μm. (d) Organoid numbers under low Wnt conditions. N = 6. * p < 0.05 compared to Mock. (e) Organoid size under low Wnt conditions. N = 6. * p < 0.05 compared to Mock. (f) Organoid numbers during long-term passages. For each passage, 1 well was split into 3 wells.

### Identification of candidates regulating Wnt-dependent epithelial regeneration in gastric organoids

To identify factors regulating Wnt-dependent regeneration in gastric epithelia, we infected gastric corpus/pyloric organoids as single cells with GeCKO A lentivirus particles, and selected reconstituted organoids with puromycin for 2 days. We then transferred the puromycin-resistant organoids to a low WNT medium (5% L-WRN+Noggin) and cultured them for 3 passages according to standard procedures described in Figure 1a and Materials and Methods. In contrast to parental pylorus/corpus organoids, which died within 2 passages under low Wnt conditions, 80 – 100 GeCKO infected organoids continued to grow after more than 3 passages per well (**Figure 1b, c)**. To identify the gene(s) whose depletion facilitated the organoid growth under low Wnt conditions, we collected surviving organoids and examined gRNA sequences inserted in their genome by direct amplicon-sequencing and TA-cloning/manual sequencing methods. We accordingly identified a list of gRNA sequences that were commonly detected in both corpus and pyloric organoids (**Figure 1d, e, f**). Importantly, gRNA’s targeting KO of *Apc*, a critical negative regulator of Wnt signalling activity were identified in the screen, validating the utility of the GeCKO screening approach for revealing modulators of Wnt-driven epithelial regeneration in the stomach. Other gRNA targets in the list identified have not previously been linked to the negative regulation of Wnt-dependent epithelial regeneration, highlighting the novelty of our screening results (**Figure 1f**).

### KO of *Alk*, *Bclaf3* and *Prkra* induces Wnt-independent stem cell maintenance

We considered genes listed in **Figure 1f** as candidate repressors of Wnt-dependent epithelial regeneration and/or activators of differentiation in gastric epithelia. Indeed, using IHC and q-PCR, we found that *anaplastic lymphoma kinase* (*Alk*), *Bclaf1 and Thrap3 family member 3* (*Bclaf3*) and *Protein activator of interferon induced protein kinase Eif2ak2* (*Prkra*) are endogenously expressed within the upper regions of mouse corpus and pyloric gastric glands, outside the *Lgr5*-expressing stem cell zone (**Supplementary Figure 3a, b**). We accordingly chose these genes for further validation.

**Figure 3.**
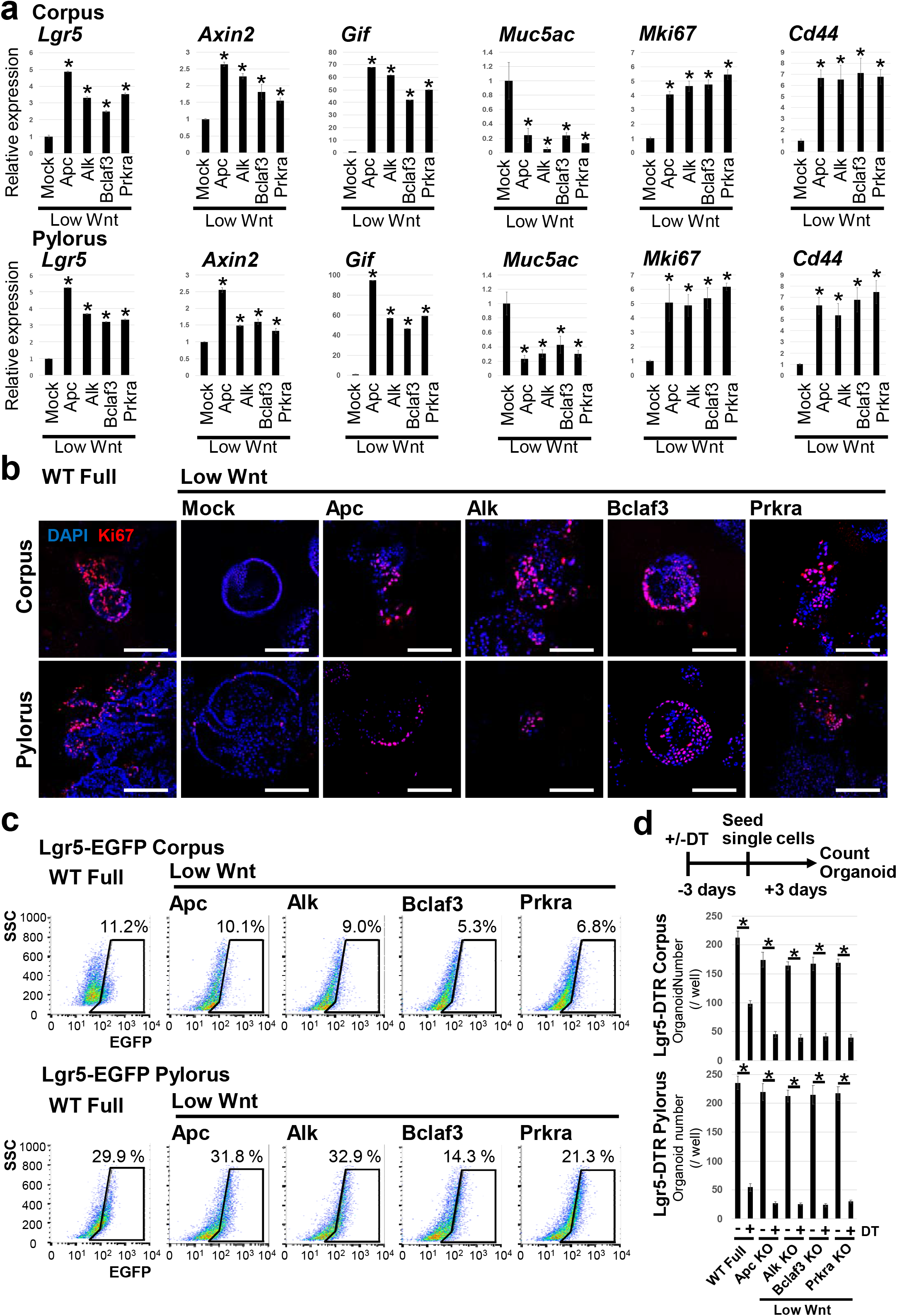
Characterization of *Alk*, *Bclaf3* and *Prkra* KO organoids growing under low Wnt conditions. (a) The expression level of lineage markers and a proliferation marker, *Mki-67*. Mock transfected organoids were cultured under low Wnt conditions for 3 days and *Apc*, *Alk*, *Bclaf3* and *Prkra* KO organoids then maintained for 3 passages under low Wnt conditions. The expression level of lineage markers was confirmed by qPCR. * p < 0.05 compared to Mock. (b) Representative immunofluorescence images of proliferation marker, MKI-67. Scale bar = 200 μm. (c) FACS analysis of Lgr5+ stem cells in organoids which were maintained under low Wnt conditions for 3 passages. (d) Results of organoid outgrowth assay combined with DT treatment. Candidate KO organoids were initially cultured under low Wnt conditions for more than 3 passages and subsequently treated with DT for 72 hours before re-seeding the organoids as single cells to evaluate their outgrowth efficiency relative to untreated controls.

To validate the GeCKO screening results, we independently knocked-out each gene using plasmid-based CRISPR/Cas9. We cloned independent sgRNAs into the pX330 plasmid, transfected them into both corpus and pyloric organoids by lipofection and subsequently selected for an ability to grow under Wnt limiting culture conditions (**Figure 2a).** After 3 passages, we confirmed the gene KO efficiency by western blotting **(Figure 2b)** and evaluated whether the single gene KO is sufficient to confer Wnt independency by quantifying the number and size of surviving organoids (**Figure 2c**). *Alk*, *Bclaf3* and *Prkra* single KO was sufficient to confer Wnt independency in both corpus/pylorus organoids (**Figure 2d, e**).

*Apc* KO abrogated exogenous Wnt dependency in both corpus and pyloric organoids. This result re-confirms that corpus and pyloric organoids critically depend on active Wnt signals for their proliferation. Of note, KO of *Axin1* or *Axin2*, known negative regulators of endogenous Wnt signalling, did not confer Wnt independency on gastric organoid cultures (**Data not shown**). Axin1 and Axin2 are functionally redundant and for the screen, we set the MOI at 0.4 to abrogate only one gene per cell. This explains why Axin1 or Axin2 were not identified as Wnt regulators in our KO-screen.

Although the number of KO organoids generated in the Wnt-depleted media was lower in comparison to the Wnt-supplemented conditions, it was comparable to that obtained following *Apc* KO. Interestingly, once organoids were formed, the surviving KO organoids grew at same rate as those cultured under Wnt-supplemented conditions, highlighting the ability of the individual gene KO to efficiently circumvent the strict dependence on exogenous Wnt ligands (**Figure 2e)**. This was further confirmed by the ability of these KO organoids to grow for more than 10 passages under low Wnt conditions (**Figure 2f)**.

To exclude any potential off-target CRISPR/Cas9 sgRNA contributions to the observed phenotype, we repeated the gene KO experiments using independently designed gRNAs (**Supplementary Figure 4a**). The independent gRNA set also induced Wnt independency in the gastric organoids, confirming the role of *Alk*, *Bclaf3* and *Prkra* in epithelial regeneration *ex vivo* (**Supplementary Figure 4b, c, d**).

**Figure 4.**
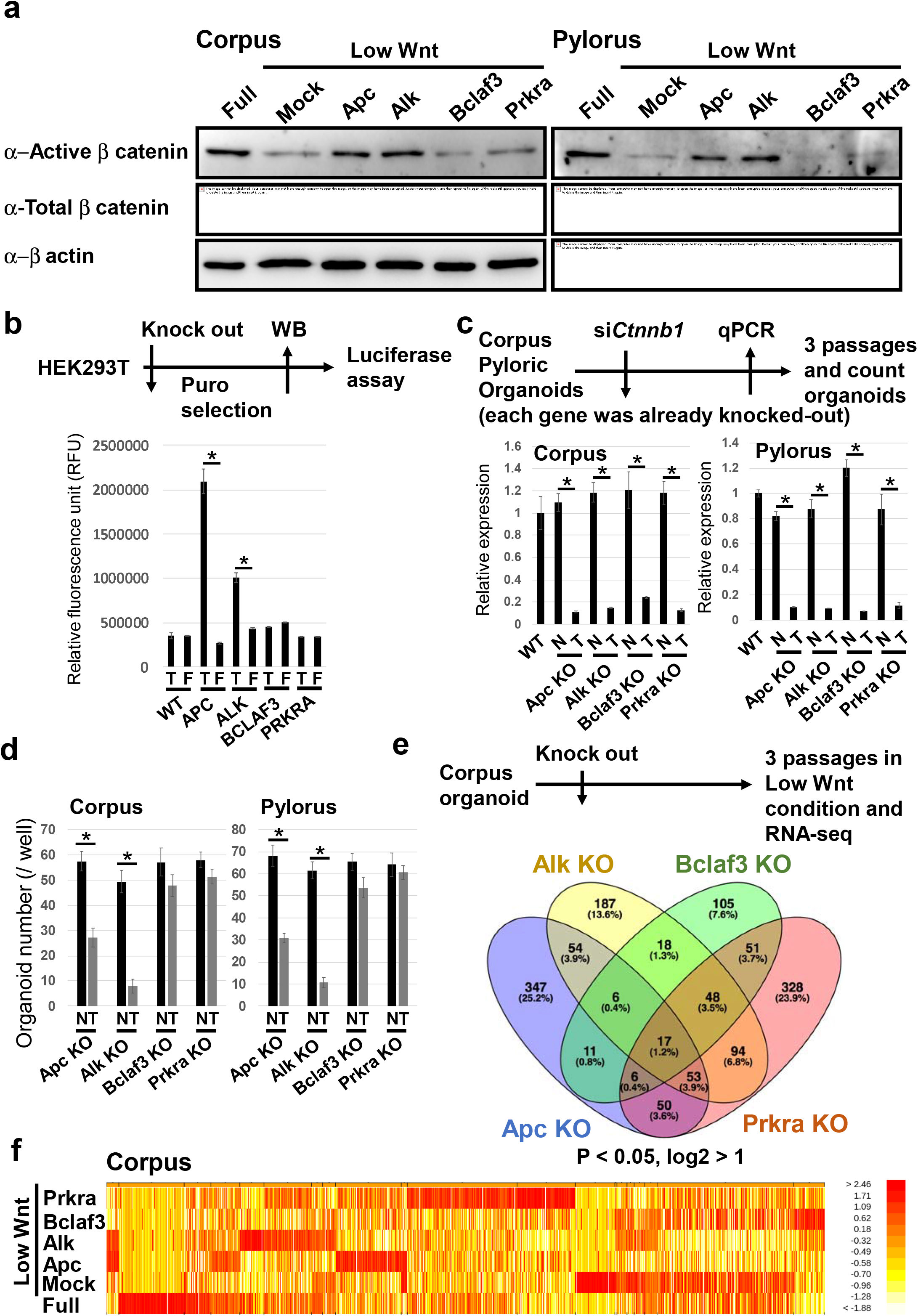
Clarification of the Wnt independency and confirmation of the downstream mechanism in *Alk*, *Bclaf3* and *Prkra* KO gastric organoids. (a) Confirmation of active β-catenin levels in each KO clone by Western Blotting. (b) TOPFlash assay using HEK293T cells. Each gene was knocked-out in HEK293T and Wnt pathway activity measured by TOPFlash assay. T; TOP signal, F; FOP signal. * p < 0.05. (c) KD efficiency of *Ctnnb1* (β-catenin) in each KO clone confirmed by qPCR. N; Non-target siRNA, T; *Ctnnb1*-target siRNA. * p < 0.05. (d) Organoid number of candidate gene K.O./*Ctnnb1* KD lines under low Wnt conditions. * p < 0.05. (e) Schematic of the RNA-seq experiment and a venn diagram which summarizes the results of RNA-sequencing. (f) A heat map showing the gene expression patterns of KO organoids. Each KO organoids show unique gene expression patterns.

Collectively, these observations strongly indicate that the single KO of *Alk*, *Bclaf3* or *Prkra* is sufficient to evade the strict dependency of gastric organoids for exogenous Wnt ligands.

To further characterize these Wnt ligand-independent organoids, we examined the expression of cell lineage, proliferation and apoptosis markers by qPCR and immunofluorescence staining. We found that KO of *Alk*, *Bclaf3* and *Prkra* increased expression of Wnt target genes and stem cell markers *Lgr5*, *Axin2* and *Gif*, whilst decreasing expression of a pit mucus cell marker, *Muc5ac* under low Wnt conditions in comparison to empty vector-transfected control organoids (**Figure 3a**). Also, KO of these genes enhanced cell proliferation, as visualized by increased *MKi67* expression and elevated expression of the stem cell marker *Cd44*, whilst decreasing apoptotic cell death (**Figure 3 a, b and Supplementary Figure 5**).

**Figure 5.**
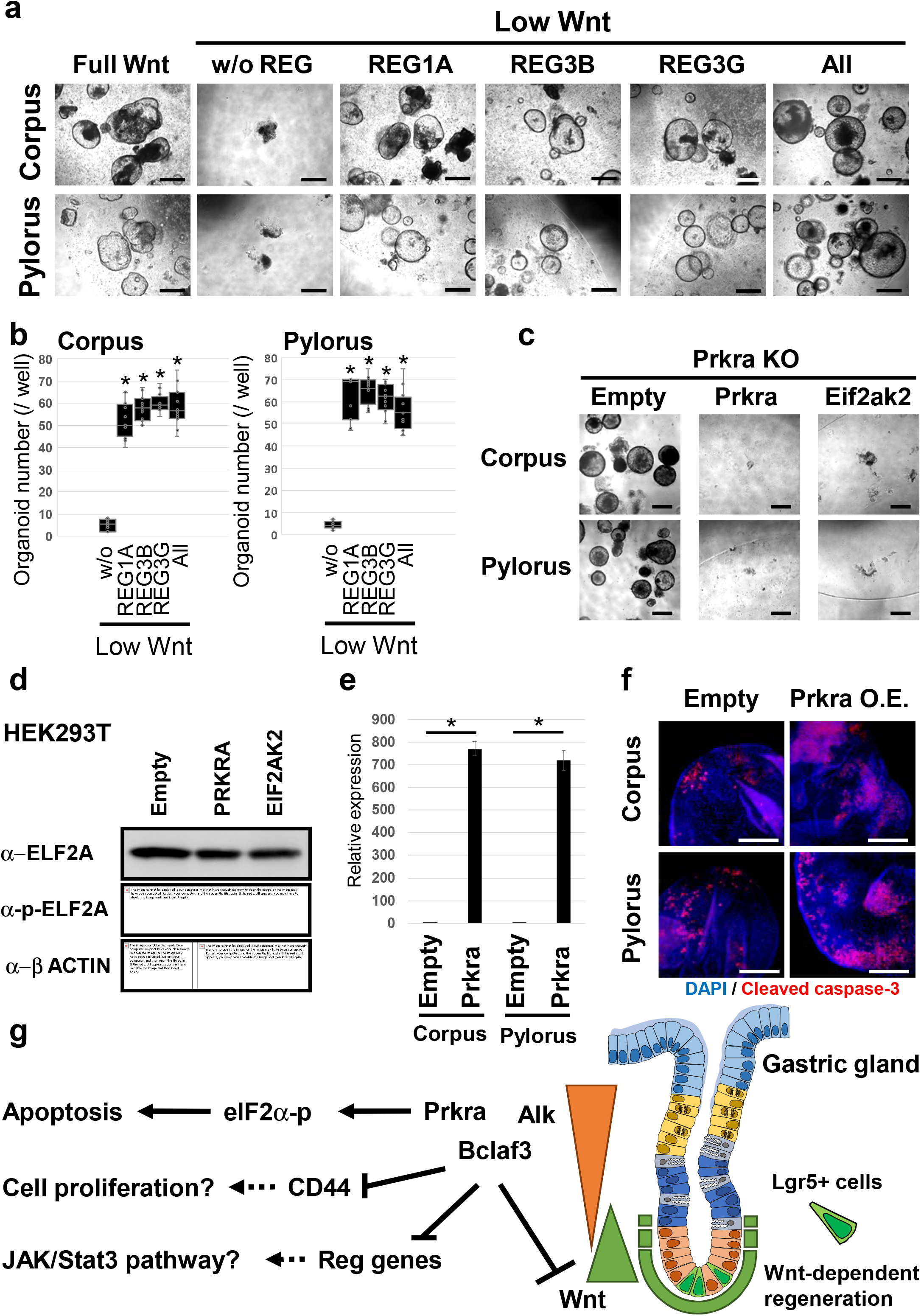
Addition of Reg proteins reduces the Wnt dependency and Prkra induces apoptosis in stomach organoids. (a) Representative images of normal gastric organoids passaged 3 times in the presence of Reg recombinant proteins. Scale bar = 200 μm. (b) Organoid numbers in the presence of Reg proteins. N = 6. * p < 0.05. (c) Increased phospho-elF2α levels following *Prkra* and *Eif2ak2* overexpression in HEK293T. OE of *Prkra* and *Eifak2* induces apoptosis in organoids, therefore we tested the effect of OE in HEK293T cells. (d) Induction of phospho-elF2α by *Prkra* and *Eif2ak2* counteract the effect of *Prkra* KO suggesting that *Prkra* KO inhibits the apoptosis induced by the Prkra-Eif2ak2-elF2α axis. Scale bar = 200 μm. (e) Confirmation of Tet-on induction of *Prkra* expression in corpus and pylorus organoids. * p < 0.05. (f) Detection of an apoptosis marker, Cleaved Caspase-3 after 24 hours of *Prkra* induction. Scale bar = 100 μm. (g) Summary cartoon. *Alk*, *Bclaf3* and *Prkra* expressed on differentiated compartments suppress Wnt signalling and expression of *Reg* genes and stem cell marker CD44. Additionally, *Prkra* regulates apoptosis. Concerted actions of these signals ensure the strict regulation of Wnt-driven, stem cell dependent epithelial regeneration in the gastric gland.

Next, we directly evaluated the effect of *Alk*, *Bclaf3* and *Prkra* KO on Lgr5^+^ stem cells via FACsorting and targeted ablation. FACs analysis of Lgr5-EGFP expression showed that KO organoids retained Lgr5^+^ gastric stem cells even under low Wnt conditions (**Figure 3c)**. To validate whether functional Lgr5^+^ stem cells are maintained in these Wnt-independent organoids, we performed organoid outgrowth assays following targeted ablation of Lgr5^+^ cells via DT treatment of Lgr5-DTRorganoid. The outgrowth efficacy from DT-treated KO organoids was dramatically reduced relative to untreated KO organoids, suggesting that *Alk*, *Bclaf3* and *Prkra* KO maintains functional Lgr5^+^ stem cells to support organoid growth even under low Wnt conditions (**Figure 3d**).

Collectively, these results indicate that KO of *Alk*, *Bclaf3* or *Prkra* supports organoid growth under Wnt limiting conditions by maintaining the resident Lgr5^+^ stem cell population.

### Deciphering underlying mechanisms of Wnt-independent cell proliferation in gastric organoids

Next, we examined the mechanism by which the novel factors regulate Wnt-driven, stem cell-dependent epithelial regeneration in gastric organoids. Firstly, to evaluate whether these factors simply function as negative regulators of Wnt signalling, we knocked-out each gene and maintained organoids in low Wnt medium for 3 passages. The activation status of the Wnt pathway was subsequently determined by quantifying activated β-catenin (Non-phospho-β-catenin) levels via western blotting. KO of *Apc* and *Alk* in both corpus and pyloric organoids increased active β-catenin levels even under low Wnt ligand conditions (**Figure 4a**). Similar modulation of Wnt activity by *APC* and *ALK* was observed via TOPFlash reporter assays following KO in HEK293T cells (**Figure 4b and Supplementary Figure 6a**). These results indicate that *Alk* is a novel negative regulator of Wnt signalling in gastric epithelial cells.

Although *Bclaf3* and *Prkra* KO organoids could also adapt to growing in medium depleted for Wnt/R-spondin (**Supplementary Figure 6b**), there was no associated increase in the level of active β-catenin in the gastric organoids, or concomitant activation of TOPFlash in HEK293T cells (**Figure 4a, b)**. This suggests that Bclaf3 and Prkra regulate epithelial regeneration independently of canonical Wnt signalling. To further substantiate this hypothesis, we additionally knocked down *Ctnnb1* in *Apc*, *Alk*, *Bclaf3* and *Prkra* KO organoids and maintained them under low Wnt conditions for 3 passages (**Figure 4c**). We reasoned that if *Bclaf3* and *Prkra* KO organoids still depend on the Wnt pathway, the KD of β-catenin should inhibit cell growth completely. As expected, *Ctnnb1* KD in *Apc* and *Alk* KO organoids robustly inhibited organoid growth, confirming their dependence on active Wnt signalling. In contrast, both *Bclaf3* and *Prkra* KO organoids continued to proliferate and expand at rates comparable to non-target siRNA transfected control organoids even under low Wnt/β-catenin conditions (**Figure 4d**). We further evaluated whether the Wnt signal inhibitors, IWR-1-endo, XAV939 and IWP-2 can suppress the proliferation of *Bclaf3* and *Prkra* KO organoids. We cultured *Bclaf3* or *Prkra* KO organoids without Wnt/R-Spondin in the presence of the Wnt inhibitors for 3 passages. In contrast to the WT/full Wnt condition, *Bclaf3* and *Prkra* KO organoids continued to proliferate and expand (**Supplementary Figure 6c**). Collectively, these results imply that KO of *Bclaf3* and *Prkra* stimulates regeneration of normal gastric organoids independently of canonical Wnt signalling.

To better define the mechanism of action of the novel factors, we performed RNA-sequencing of *Apc*, *Alk*, *Bclaf3* and *Prkra* KO organoids. We generated *Apc*, *Alk*, *Bclaf3* and *Prkra* KO organoids via the plasmid-based CRISPR/Cas9 system and selected clones under low Wnt conditions for 3 passages before performing RNA-sequencing (**Figure 4e**). Although each KO sample showed a unique gene expression profile (**Figure 4e, f**), we found that a set of genes and pathways (e.g. Cadherin signalling, Inflammation mediated chemokine and cytokine signalling) were commonly upregulated in those KO organoids **(Supplementary Figure 7a).** Importantly, some of the genes identified in the Lgr5^+^ stem cell transcriptome were also upregulated in *Alk*, *Bclaf3* and *Prkra* KO organoids **(Supplementary Figure 7b).** Therefore, those genes and pathways may be important for the maintenance of epithelial stem cells, and endogenous *Alk*, Bclaf3 and *Prkra* could be instrumental in directing cell differentiation by suppressing those genes and pathways.

### *Reg* family genes promote cell proliferation under Wnt-limiting conditions

Among the commonly up-regulated genes, we focused on *regenerating islet-derived (Reg)* family genes, which encode small secretory proteins. *Reg* gene expression was significantly increased in the KO organoids compared to wild-type organoids when cultured for 3 days under Wnt-limiting conditions (**Supplementary Figure 7b, c**). Additionally, endogenous *Reg* gene expression was strongest at the base of gastric glands *in vivo*, where endogenous stem cells reside (**Supplementary Figure 8a, b**). Thus, we examined whether up-regulation of *Reg* gene family expression contributes to the sustenance of KO organoid growth under Wnt-limiting conditions. We cultured normal corpus and pyloric organoids for 3 passages in low Wnt/R-spondin medium supplemented with individual recombinant REG proteins (**Figure 5a**). In contrast to the lack of organoid growth in low Wnt/R-Spondin media, both pyloric and corpus organoids continued to expand in REG protein-supplemented media (**Figure 5b**). Notably, concomitant addition of the various recombinant REGs failed to further enhance organoid growth, suggesting that *Reg1*, *Reg3β* and *Reg3γ* are functionally redundant for the growth of epithelial cells.

Similar positive effects on epithelial proliferation were observed following overexpression of *Reg* family genes in gastric organoids (**Supplementary Figure 9a, b, c, Supplementary Figure 10a**). Those organoids could be maintained for at least 5 passages under low Wnt/R-Spondin conditions (**Supplementary Figure 9d**). Of note, elevated Reg activity leads to increased expression of the epithelial stem cell marker gene *Lgr5* and proliferation markers such as *Mki67* (**Supplementary Figure 10b**). Therefore, Reg proteins may be novel regulators of epithelial regeneration through their ability to partially substitute for canonical Wnt signalling and promote expansion of the Lgr5^+^ stem cell compartment.

### *Prkra* KO blocks cell death by limiting phosphorylation of elf2alpha proteins

In gastric epithelia, apoptotic cell death is observed within the upper region of gastric glands under homeostatic conditions (20). It has been reported that Prkra induces the phosphorylation of elf2α, leading to cell death (21). We therefore reasoned that KO of *Prkra* might promote epithelial expansion *ex vivo* through suppression of cell death via inhibition of endogenous elF2α phosphorylation. To examine this possibility, we over-expressed *Prkra* and *Eif2ak2*, a serine/threonine-protein kinase which transmits stress signals by phosphorylated elF2α in *Prkra* KO gastric organoids, and cultured them for 3 passages under Wnt-limiting conditions. We showed that over-expression of *Prkra* and *Eif2ak2* counteracted the organoid growth observed following *Prkra* KO (**Figure 5c)**. Since constitutive over-expression of *Prkra* and *Eif2ak2* via PiggyBac vectors induced rapid cell death of normal gastric organoids, we over-expressed *PRKRA* and *EIF2AK2* in HEK293T cells and confirmed associated phosphorylation of ELF2A (**Figure 5d**). Furthermore, to test whether *Prkra* directly induces apoptosis, we temporally induced *Prkra* and stained those organoids with the apoptosis marker, Cleaved caspase-3. As a result, we confirmed that induction of *Prkra* increased apoptosis in normal gastric organoids (**Figure 5e, f**). These results suggest that *Prkra* KO can support epithelial regeneration by suppressing apoptotic cell death via limiting phospo-elF2α levels.

Our data collectively reveal that *Alk*, *Bclaf3* and *Prkra* function to suppress Wnt-related and *Reg* family genes and enhance apoptosis to regulate epithelial regeneration in the stomach. These observations readily explain why KO of these genes enhances epithelial expansion independently of Wnt signalling in our organoid screen (**Figure 5 g)**.

## DISCUSSION

Wnt signalling is of critical importance for maintaining the integrity of gastric epithelia via regulation of endogenous stem cell function (5). In mouse gastric epithelia, resident Lgr5-expressing populations at the gland base in the pylorus and corpus function as homeostatic and injury-activated stem cell populations respectively (3, 4). *Lgr5*, which is itself a Wnt target gene, functions as a facultative component of the Wnt receptor complex, where it is thought to facilitate binding of the Wnt agonist, R-spondin, to modulate Wnt signalling levels activated by Wnt ligands provided by surrounding mesenchyme to a level compatible with optimal stem cell function (22). However, detailed mechanistic insight into the regulation of Wnt signalling and the delicate balance between self-renewal and differentiation in the stomach is currently lacking. To try and address this knowledge gap, we established a new Genome-Scale CRISPR knock-Out screening technique on normal gastric organoids and identified novel factors which potentially regulate either endogenous Wnt signalling or cell differentiation in mouse gastric epithelia. We found that KO of *Alk*, *Bclaf3* and *Prkra* supports Wnt-independent regeneration and apoptosis in gastric epithelia, whilst concomitantly inducing expansion of the Lgr5^+^ stem cell population. Taken together with the observation that these factors are predominantly expressed outside the Lgr5^+^ stem cell zone in mouse glandular epithelia, this indicates they may function to suppress stemness and maintain/promote the differentiation status of gastric epithelial cells. Mechanistically, these factors suppress Wnt signalling, Integrin signalling and Inflammation mediated by the chemokine and cytokine signalling pathways. Of note, those factors may suppress *Reg* family genes, which support stemness in gastric epithelia. Although further verification is needed, results here suggest that *Alk*, *Bclaf3* and *Prkra* may function as novel regulators of gastric epithelial differentiation (**Figure 5g**).

We show that *Alk* is predominantly expressed outside the Lgr5^+^ stem cell zone and functions to suppress endogenous Wnt signalling. A similar result was observed in HEK293T cells, indicating that *Alk* may be a novel Wnt antagonist in different tissues. In cancer, *Alk* has been reported as a common fusion partner for other genes to promote tumour malignancy (23), although it is not known whether this involves modulation of the Wnt pathway. Our results indicate that *Alk* itself can suppress Wnt signalling in *ex vivo* gastric organoids, potentially as a mechanism to ensure terminal differentiation towards the various functional lineages that populate the epithelial glands.

Our data also implicates *Bclaf3* and *Prkra* as novel regulators of gastric epithelial growth. Some Wnt ligands were upregulated by *Bclaf3* and *Prkra* KO, resulting in the Wnt signalling pathway being identified as one of the top up-regulated pathways (**Supplementary Figure 7b**). However, our IWP2 treatment results (**Supplementary Figure 6c**) indicate that these upregulated Wnts may not be suffficient to maintain gastric regeneration. It has been reported that Bclaf3 binds to the Oct-Sox binding motif and is necessary to maintain pluripotency of mouse embryonic stem cells (mESCs) (24). In contrast to this report, we find that *Bclaf3* expression is present in non-stem cells and may induce differentiation in gastric glands by supressing Wnt signalling and *Reg* family genes. Importantly, our results suggest *Bclaf3* and *Prkra* likely regulate epithelial differentiation by controlling expression of members of the *Reg* gene family, which are known to be potent modulators of epithelial regeneration and stem cell maintenance.

It has been reported that Reg expression is controlled by Wnt signalling in mESCs (25). Furthermore, Reg expression is regulated by Gastrin and also IL-6, IL-22 in human gastric cancer lines (26, 27). Mechanistically, the gp130 transmembrane protein is thought to be a Reg receptor and the gp130-JAK2-STAT3 axis promotes cell growth and resistance to apoptosis, as well as inducing poorly differentiated cells in pancreatic cancer cells (28). Indeed, Reg has a growth-promoting effect and may play an important role in the reconstitution of gastric mucosa. (20). Therefore, there is a possibility that Wnt and/or Gastrin – Regs - JAK/Stat3 signalling cascade is essential for the regeneration of gastric epithelia. Consistent with this, *Reg* KO mice shows significantly decelerated growth and migration of intestinal epithelial cells towards the villus tip (29). Indeed, several *Reg* family genes are expressed in the endogenous gastric stem cell compartment (**Supplementary Figure 8a, b)** (4, 30). Whilst administration of REGs cannot fully compensate for Wnt withdrawal, *Reg* genes clearly supported organoid growth. Supplementation of gastric organoid media with all three recombinant REGs failed to show a synergistic effect on organoid growth. This result suggests that the Reg family gene displays functional redundancy during epithelial regeneration.

Our data, together with the mutually exclusive expression pattern of *Bclaf3*/*Prkra* and *Reg* genes within the non-stem cell and stem cell compartments respectively of mouse gastric glands *in vivo*, supports a model in which the absence of *Bclaf3*/*Prkra* expression at the gland base facilitates robust *Reg* gene expression to support Lgr5^+^ stem cell function. Expression of *Bclaf3*/*Prkra* outside the stem cell zone may help facilitate terminal differentiation towards the specialized lineages of the gastric epithelia. The role of REG family genes in gastric epithelia warrants further examination in future.

Although conditional KO of *Alk*, *Bclaf3* and *Prkra* in the stomach has not been reported, conventional KO of each gene does not generate obvious gastric phenotypes. Individual KO of those genes in gastric organoids up-regulates common pathways (**Supplementary Figure 7a, b**), suggesting that *Alk*, *Bclaf3* and *Prkra* likely exhibit some functional redundancy in the stomach. One of those pathways may be upregulation of CD44. CD44 is a non-kinase transmembrane glycoprotein thought to be induced in several types of stem cells including cancer-initiating cells (REFS). CD44 activates cell proliferation, survival and modulates cytoskeletal changes to enhance cellular motility (31). Therefore, CD44 may also have a role in regulating ? stemness in the normal gastric gland.

Another pathway may be the suppression of apoptosis. It has been reported that *Prkra* induces apoptosis via phosphorylation of elF2α in both human and mouse cell lines (32, 33). Furthermore, Prkra is expressed in colonic epithelial cells, where it activates apoptosis (34). Here, we confirm that *Prkra* induces apoptosis in *ex vivo* gastric organoids (**Figure 5e, f**). Although massive induction of *Prkra* caused apoptosis, endogenous expression of *Prkra* in the gastric epithelium ??was modest (**Supplementary Figure 3a**). Given that *Alk* and *Bclaf3* may also induce apoptosis, these factors may work cooperatively to balance epithelial regeneration/cell death *in vivo*.

Our genome-wide organoid screen has revealed novel factors involved in regulating Wnt signalling, *Reg* gene expression and proliferation/apoptosis in the mouse gastric epithelium. Future studies will be directed at further refining the role of these factors in regulating the balance between self-renewal and differentiation in healthy gastric epithelium, and the potential contributions of these factors to gastric cancer. Given that healthy/cancer organoids can now be readily generated from multiple tissues of mouse and human origin, the novel screening procedure detailed should prove useful in deriving clinically-relevant biological insights across a range of healthy and disease settings.

## AVAILABILITY

CHOPCHOP is a web tool for selecting target sites for CRISPR/Cas9, CRISPR/Cpf1, CRISPR/Cas13 or NICKASE/TALEN-directed mutagenesis (https://chopchop.cbu.uib.no)

## ACCESSION NUMBERS

Amplicon sequencing data and RNA sequencing data that support the findings of this study have been deposited in NCBI Sequence Read Archive (SRA) accession code SRR12182373 - 79 and SRR12214478 – 84 respectively. All other data supporting the findings of this study are available from the corresponding author(s) on reasonable request.

## SUPPLEMENTARY DATA

Supplementary Data are available online.

## ACKNOWLEDGEMENT

The authors thank Dr. Fred de Sauvage (Genentech) for the Lgr5-DTR-EGFP mice, Dr. Hitoshi Niwa (Kumamoto University) for PiggyBac vectors, Dr. Feng Zhang (Broad Institute of MIT and Harvard) for the Mouse GeCKOv2 CRISPR knockout pooled library and the pX330-U6-Chimeric_BB-CBh-hSpCas9 plasmid, Dr. Bob Weinberg (Whitehead Institute for Biomedical Research) for pCMV-VSV-G and pCMV-dR8.2 dvpr plasmids, Dr. Randall Moon (University of Washington School of Medicine) for M50 Super 8x TOPFlash and M51 Super 8x FOPFlash plasmids. The authors thank Ms. Ayako Tsuda and Ms. Manami Watanabe for technical assistance.

## AUTHOR CONTRIBUTION

K.M. designed, performed experiments, collected, analysed data and wrote the manuscript. Y.T., H.T. designed, performed experiments and collected data. K.S. and Y.J. provided technical help. M.O. analysed data and provided advice. N.B. supervised the project, analysed data and wrote the manuscript. All authors discussed results and edited the manuscript.

## FUNDING

K.M. is supported by Japan Society for the Promotion of Science [grant number 17K07161]. Y.T is supported by Japan Society for the Promotion of Science [grant number 17H06710]. H.T. is supported by Japan Society for the Promotion of Science [17H03586]. N.B. is supported by the Agency for Science, technology and Research (A*STAR); the National Research Foundation, Prime Minister’s Office,   510 Singapore under its Investigatorship Program (Award No. NRF-NRF12017-03) and Japan Society for the Promotion of Science [grant number 17H01399]. Funding for open access charge: Japan Society for the Promotion of Science [grant number 17H01399]

## CONFLICT OF INTEREST

The authors declare no competing interests.

## Notes

### Competing Interest Statement

The authors have declared no competing interest.

