## Supplementary figures for "Genome-Scale CRISPR Screening reveals novel factors regulating Wnt-dependent regeneration of mouse gastric organoids"

#### Supplementary Figure1_KMurakami

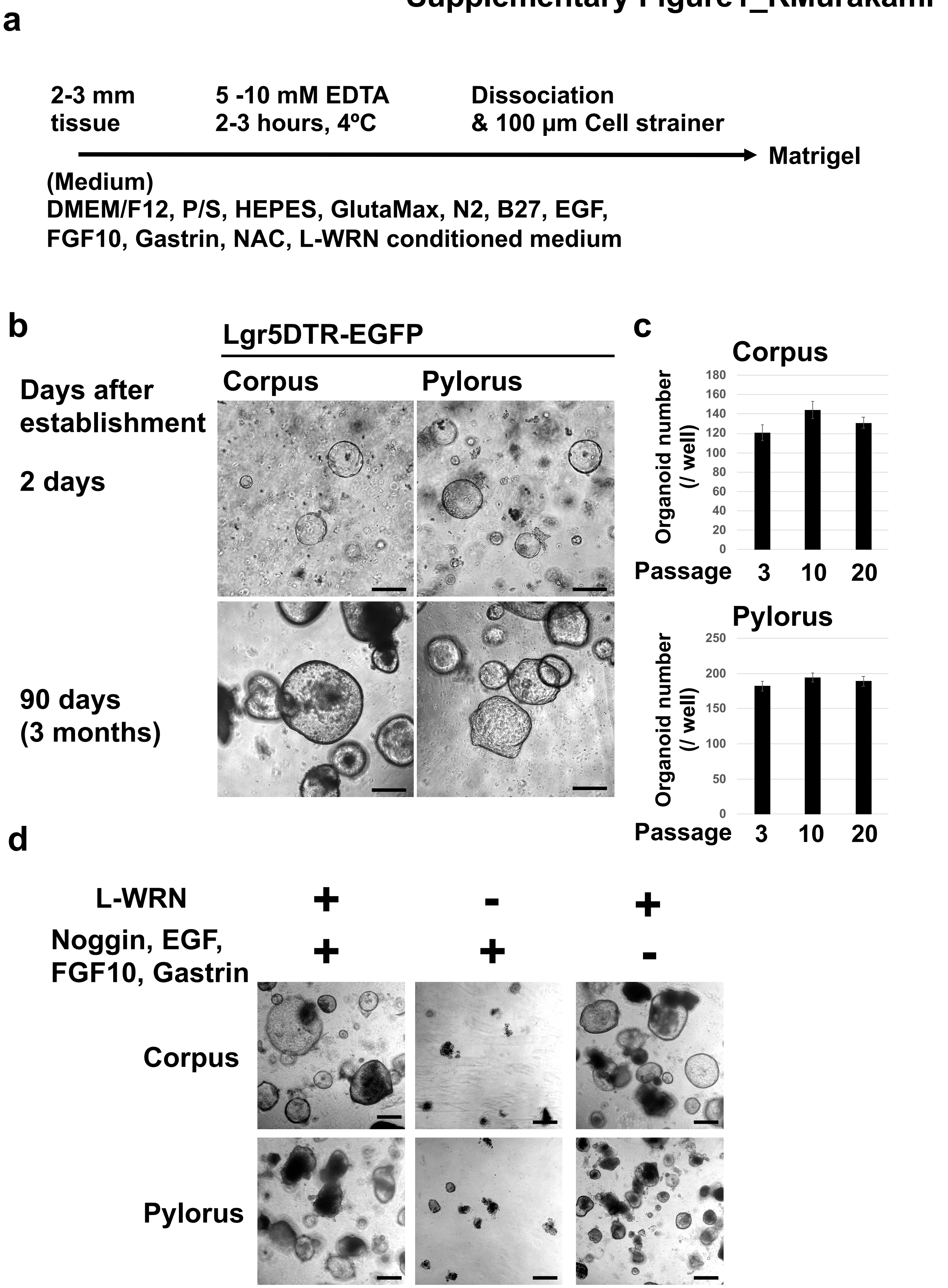


**Supplementary Figure 1.** (a) Procedure for the establishment of mouse stomach organoids. (b) Images of organoids after establishment and following passage for 3 months. Scale bar = 200 µm. (c) Organoid outgrowth efficiency over multiple passages. Organoids were dissociated and single cells were seeded in Matrigel. After 3 days, the number of reconstituted organoid was counted. N = 3. (d) Confirmation of Wnt signal dependency of stomach organoids. Scale bar = 200 µm.

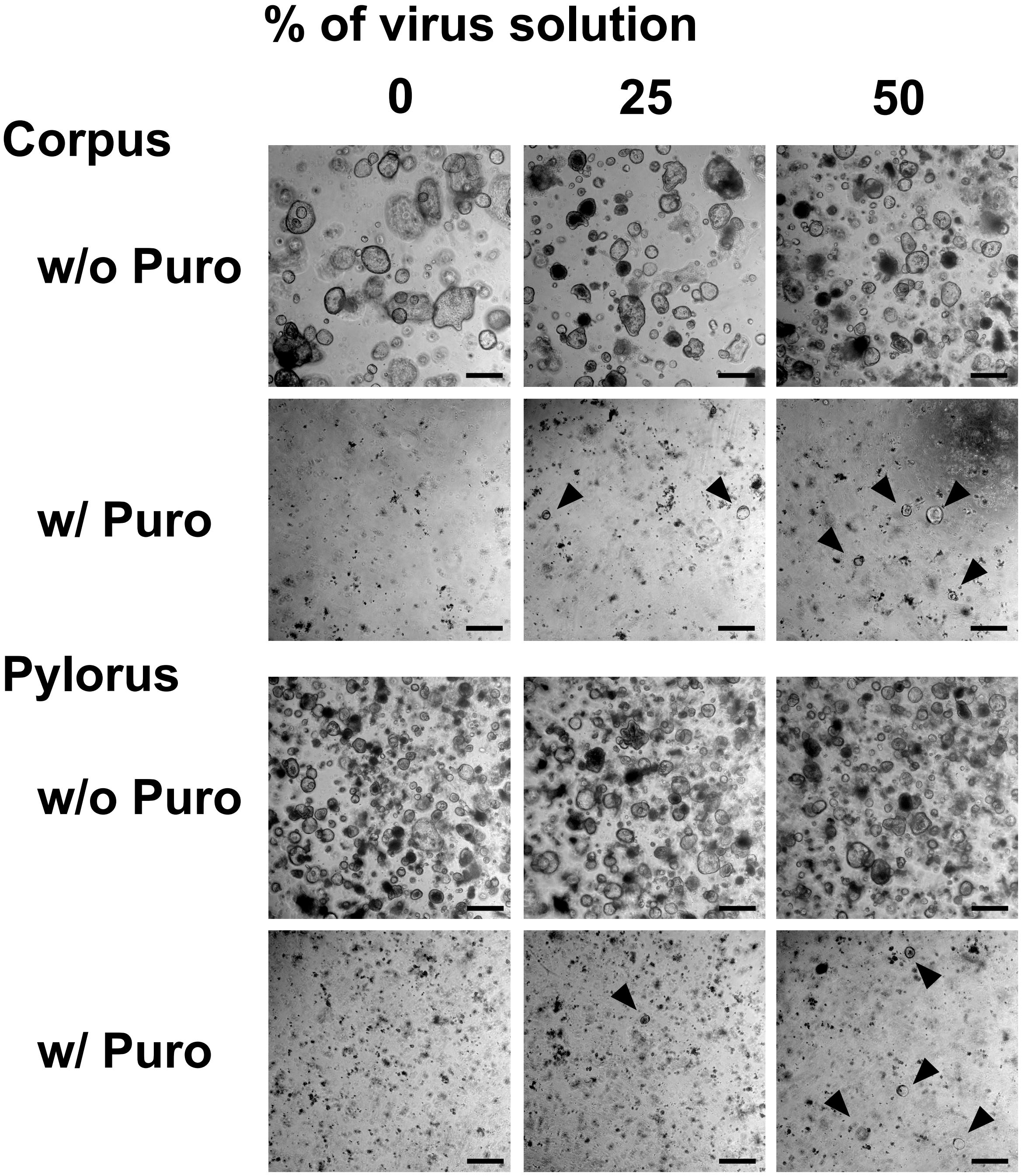


**Supplementary Figure 2.** Confirmation of MOI for the GeCKO screening. Arrowhead; Reconstituted Puro resistant organoids. Scale bar = 200 µm. See Materials and Methods for the details.

Supplementary Figure3_KMurakami

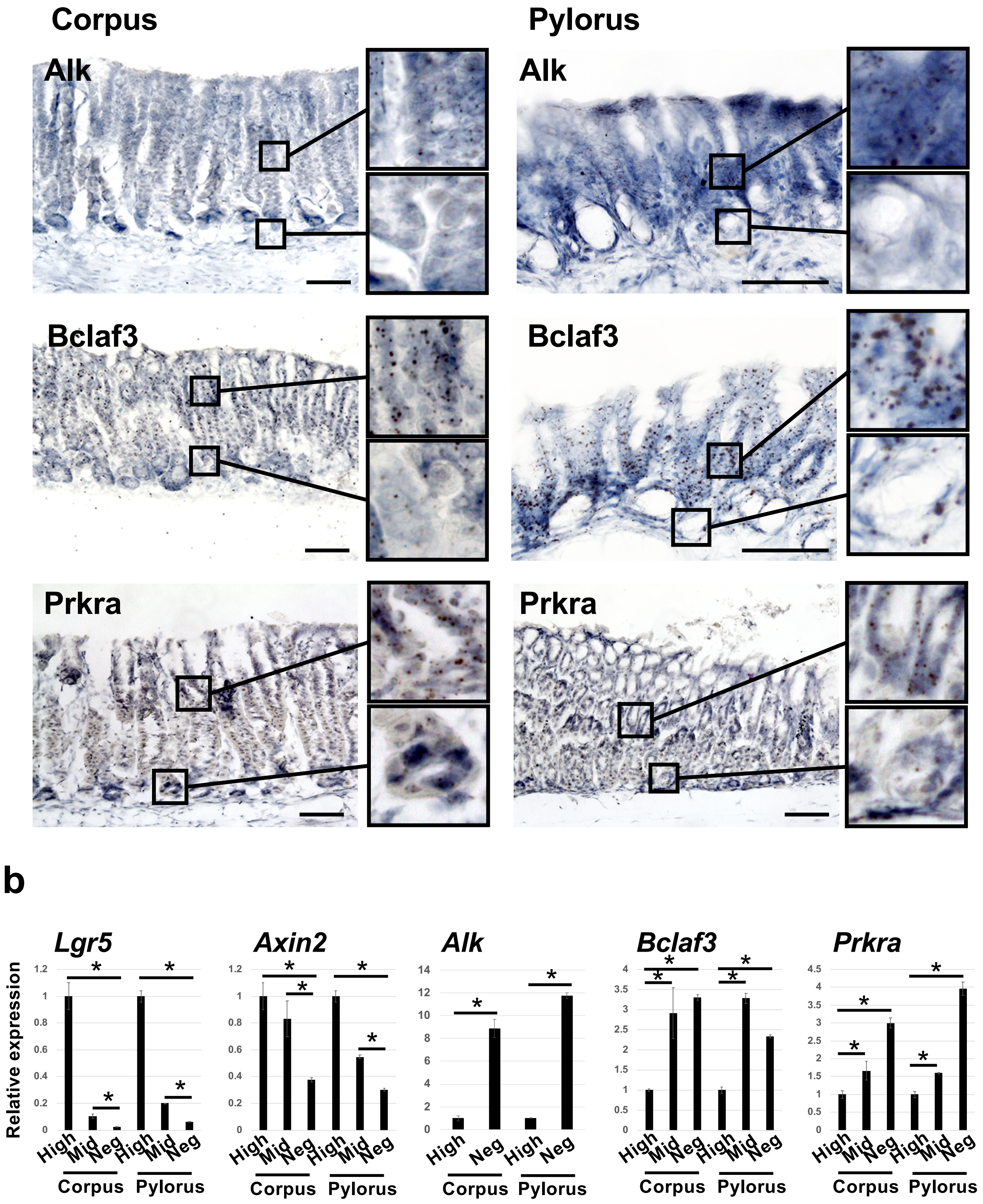


**Supplementary Figure 3.** (a) Expression status of *Alk*, *Bclaf3* and *Prkra in vivo* mouse stomach. mRNA expression was confirmed by RNA *in Situ* hybridization. Scale bar = 50 μm. (b) Expression of *Alk*, *Bclaf3* and *Prkra* in Lgr5 high/ middle/ negative cells sorted from *in vivo* mouse stomach. The mRNA expression level was confirmed by qPCR. * $p < 0.05$.

#### Supplementary Figure4_KMurakami

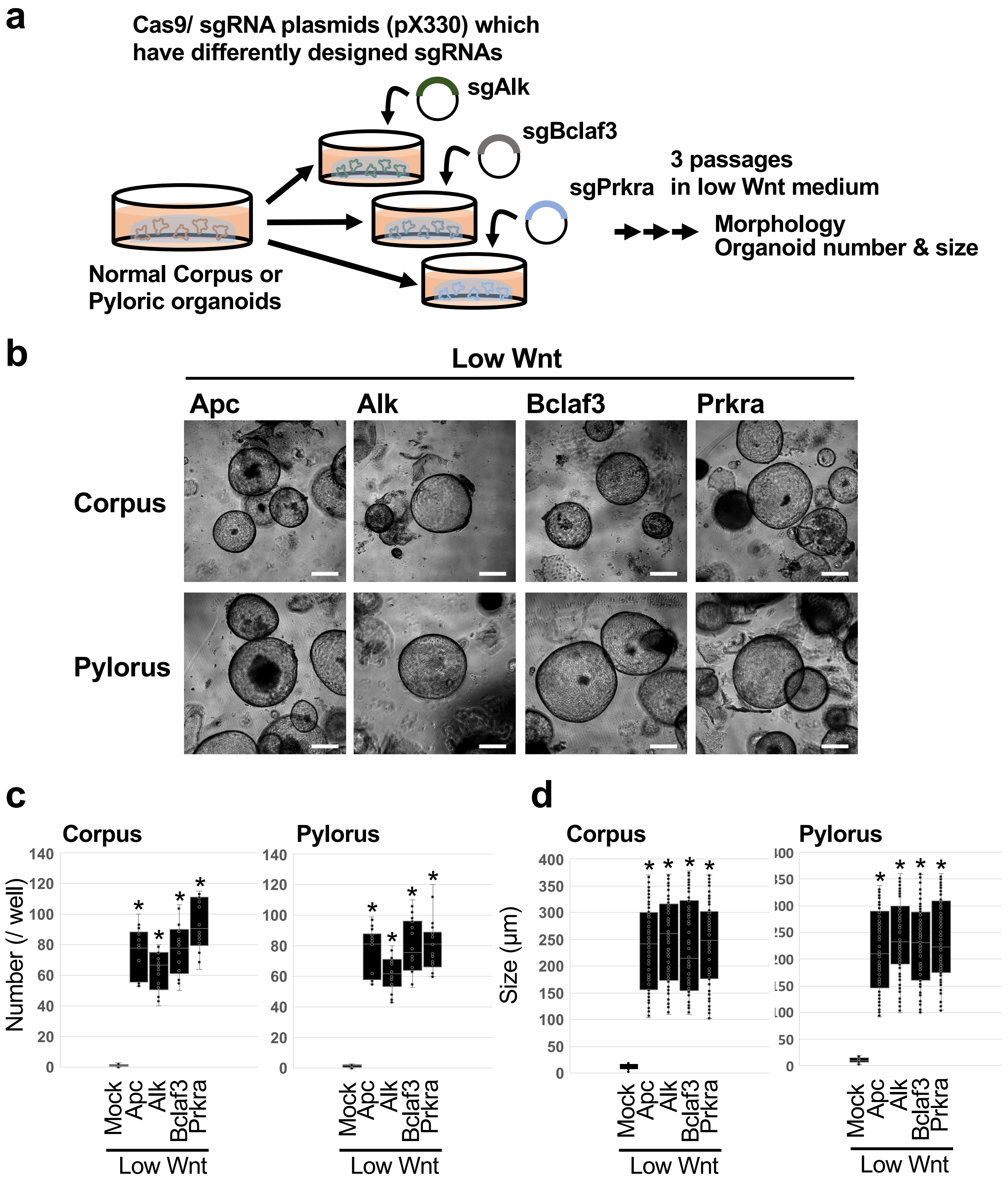


**Supplementary Figure 4.** (a) Validation strategy: Newly designed single sgRNAs were cloned into the CRISPR/Cas9 plasmid (pX330) and transduced into stomach organoids. Organoids were selected under low Wnt conditions for 3 passages. (b) Representative images of single-gene knockout organoids after 3 passages under low Wnt conditions. Scale bar = 200 µm. (c) The organoid number under low Wnt conditions. N = 6. * $p < 0.05$ compared to Mock. (d) Organoid size under low Wnt conditions. N = 6. * $p < 0.05$ compared to Mock.

### Supplementary Figure5_KMurakami

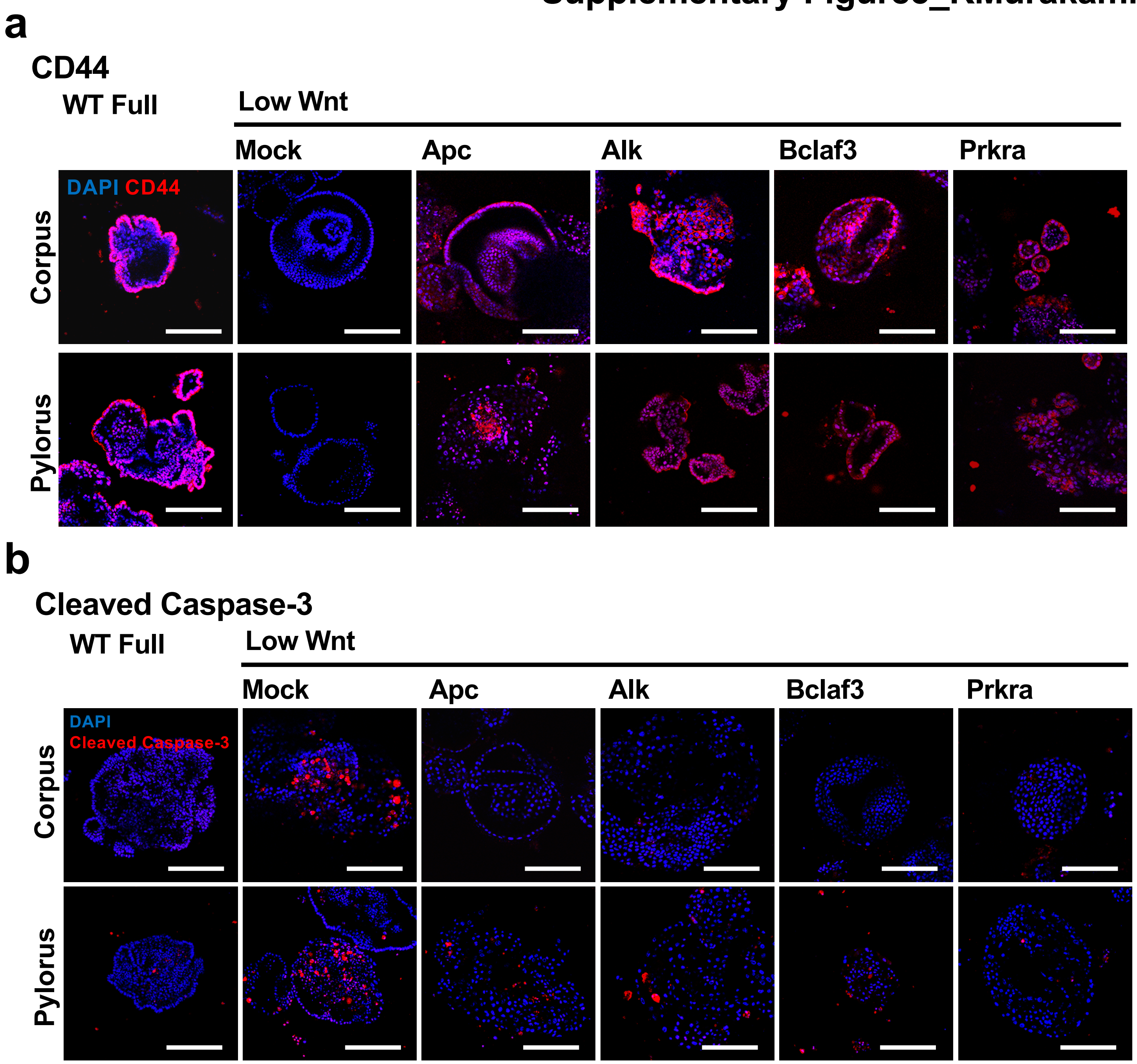


**Supplementary Figure 5.** (a) Representative immunofluorescence images of a stem cell marker, CD44 after 3 passage under low Wnt conditions. Scale bar = 200 µm. (b) Representative immunofluorescence images of the apoptosis marker Cleaved caspase-3 after 3 passage under low Wnt conditions. Scale bar = 200 µm.

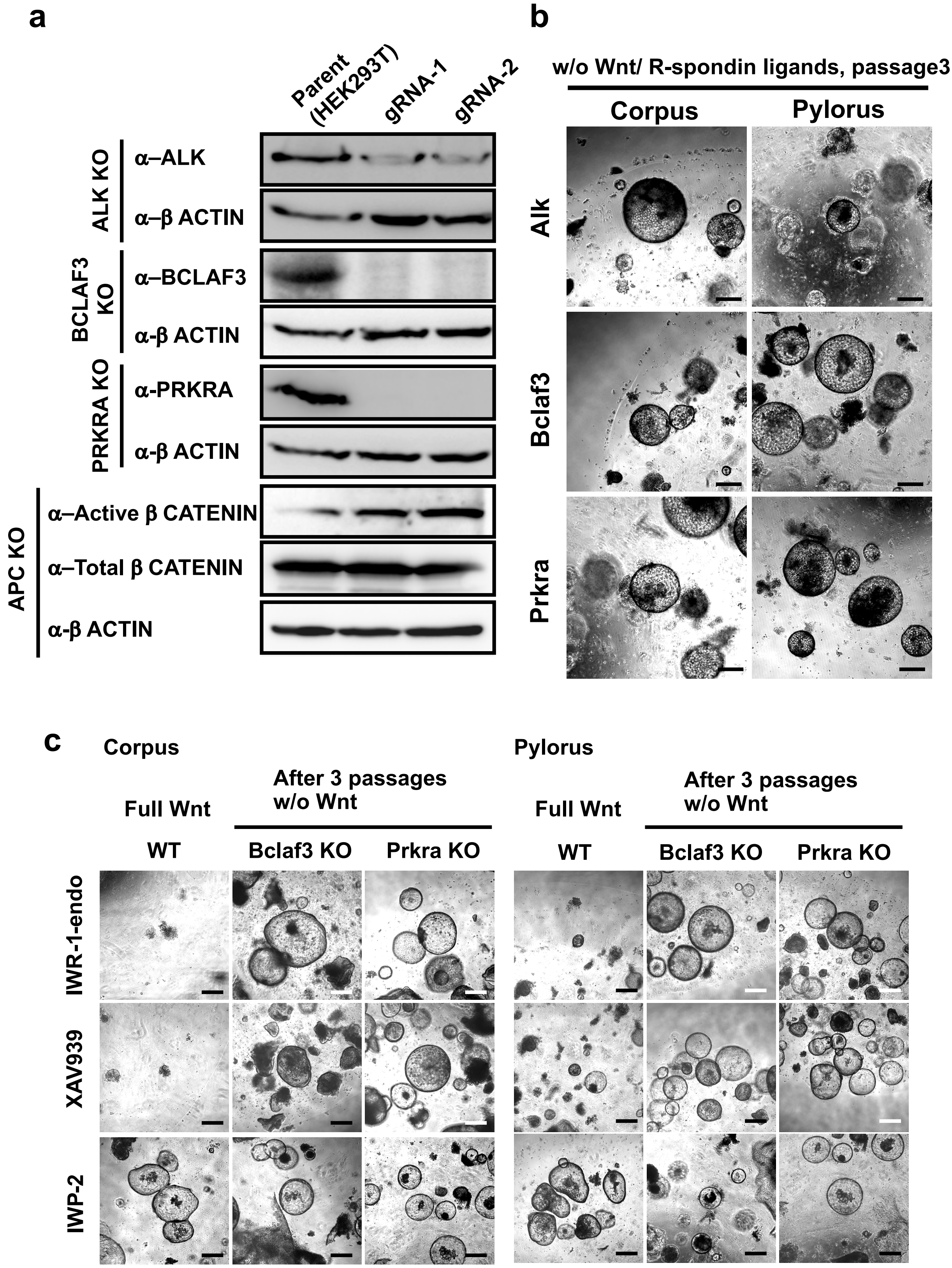


**Supplementary Figure 6.** (a) Confirmation of gene KO in HEK293 cells by Western Blotting. pX330 plasmids, which express independent gRNAs were transfected with a CAG-Puro$^r$ plasmid into HEK293T cells by lipofection. After a brief selection of puromycin, knock-out efficiency was confirmed by Western Blotting. For APC KO, we confirmed the activation of β-catenin instead of APC protein itself. (b) Representative images of organoids grown in the absence of Wnt/R-spondin ligands. Scale bar = 200 μm. (c) Representative images of organoids grown under in the absence of Wnt activation. KO organoids were passaged for 3 times in the presence of Wnt inhibitors. IWR-1-endo and XAV939, tankyrase inhibitors, stabilize Axin and enhance the degradation of β-catenin, resulting in the inhibition of the Wnt pathway. IWP-2, a porcupine inhibitor inhibits the secretion of Wnt ligands. Scale bar = 200 μm.

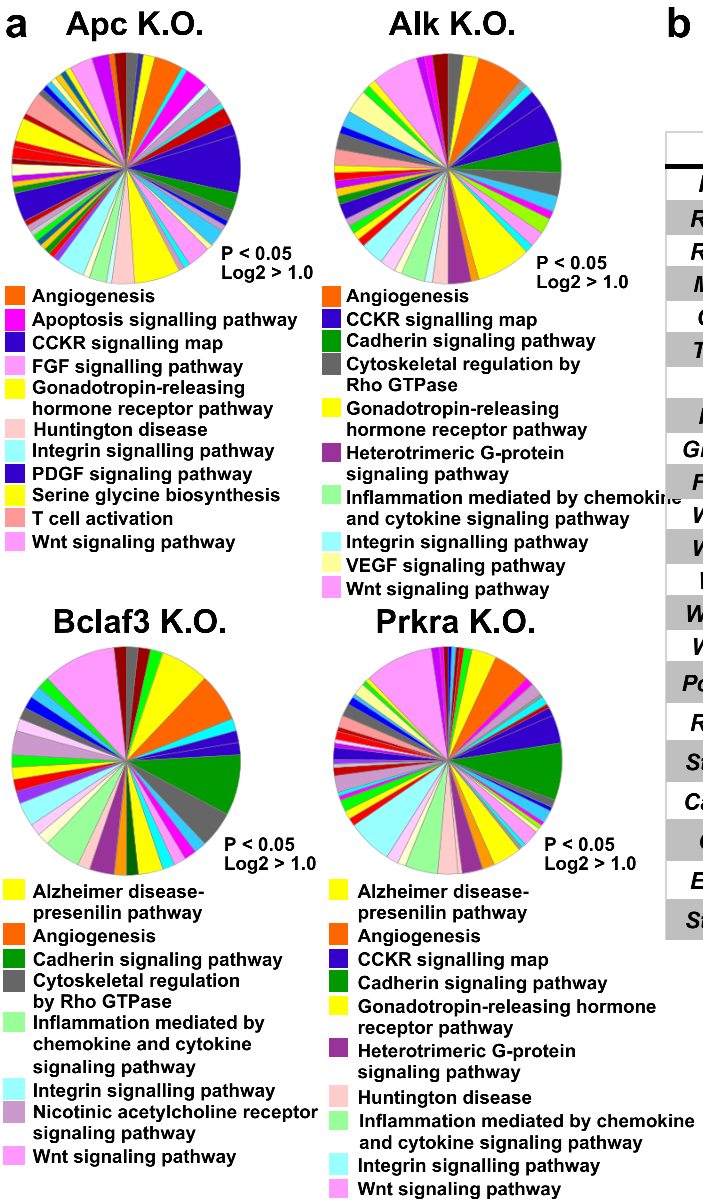


**b**

**Log2 fold changes**

| | Alk | Bclaf3 | Prkra |
|---|---|---|---|
| *Reg1* | 3.1 | 1.1 | 5.23 |
| *Reg3b* | 4.4 | N.D. | 10.42 |
| *Reg3g* | N.D. | N.D. | 2.34 |
| *Mmp7* | N.D. | N.D. | 2.17 |
| *CD44* | 2.06 | N.D. | N.D. |
| *Thbs1* | 1.91 | 1.12 | N.D. |
| *Car8* | 10.94 | 6.29 | 7.23 |
| *Rec8* | N.D. | N.D. | 1.28 |
| *Gm6744* | N.D. | 4.29 | N.D. |
| *Fcamr* | N.D. | 1.43 | N.D. |
| *Wnt9a* | 7.02 | 6.09 | 7.7 |
| *Wnt16* | 4.56 | N.D. | 5.04 |
| *Wnt6* | 4.4 | N.D. | 5.03 |
| *Wnt10a* | 2.94 | N.D. | N.D. |
| *Wnt5a* | N.D. | 2.46 | 4.17 |
| *Pcolce2* | -1.93 | -2.05 | -1.1 |
| *Robo2* | -1.47 | -1.59 | N.D. |
| *St6gal2* | -3.05 | -3.18 | -3.44 |
| *Cacnb4* | -4.27 | N.D. | -4.66 |
| *Cpvl* | N.D. | -5.84 | N.D. |
| *Ero1lb* | N.D. | N.D. | -1.37 |
| *St3gal6* | N.D. | N.D. | -1.18 |

N.D.; No significant difference

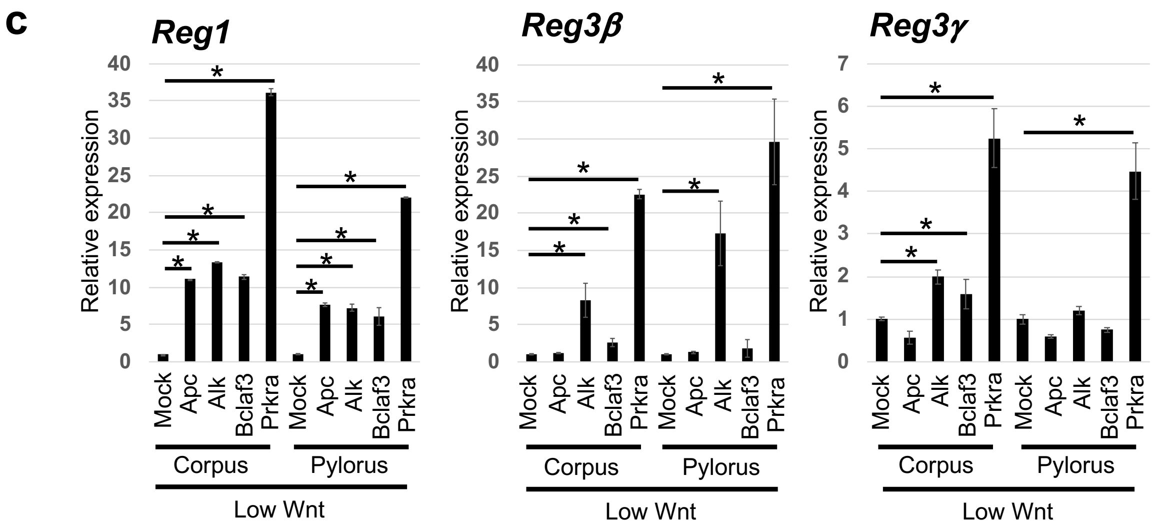


**Supplementary Figure 7.** (a) Significantly upregulated genes ($p < 0.05$, Log2 > 1) were selected from RNA-sequencing results of *Alk*, *Bclaf3* and *Prkra* knockout organoids. The resulting gene lists were further analysed by PANTHER software. Numerous pathways, including Wnt signalling are upregulated by *Alk*, *Bclaf3* and *Prkra* KO in corpus organoids. (b) Some genes preferentially expressed in Lgr5+ corpus reserve stem cells were upregulated in *Alk*, *Bclaf3* and *Prkra* KO corpus organoids. (c) Confirmation of the upregulation of *Reg1*, *Reg3b* and *Reg3g* genes in *Alk*, *Bclaf3* and *Prkra* KO organoids by qPCR. * $p < 0.05$.

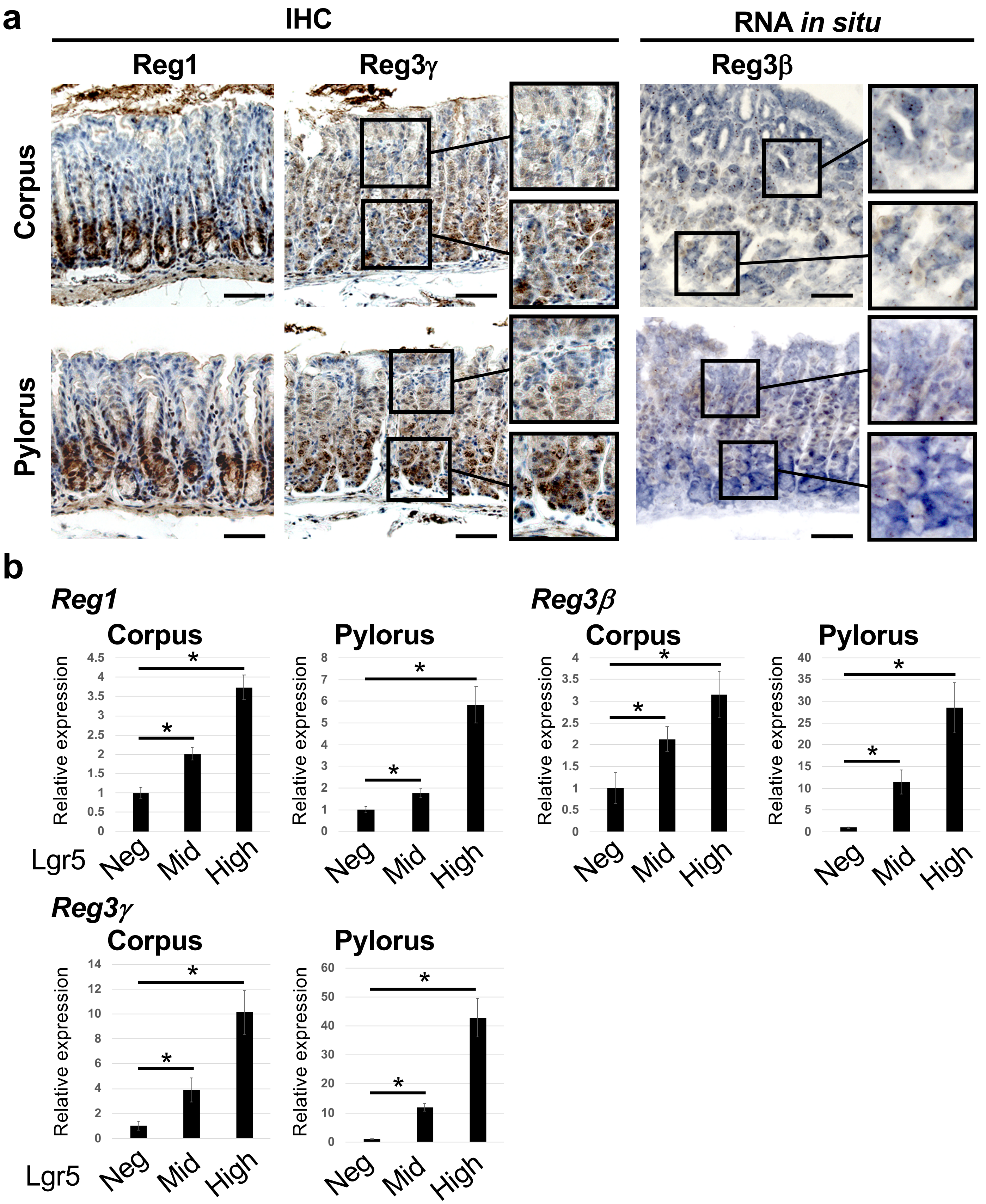


**Supplementary Figure 8.** (a) Confirmation of the expression of *Reg* family genes in stomach glands. Reg1 and Reg3γ proteins were visualized by immunostaining and Reg3β mRNA was detected by RNA *In Situ* hybridization. Scale bar = 50 µm. (b) Confirmation of Reg1, Reb3β and Reg3γ expression in Lgr5 high/ middle/ negative cells collected from *in vivo* mouse normal stomach. mRNA expression level was confirmed by qPCR. * $p < 0.05$.

### Supplementary Figure9_KMurakami

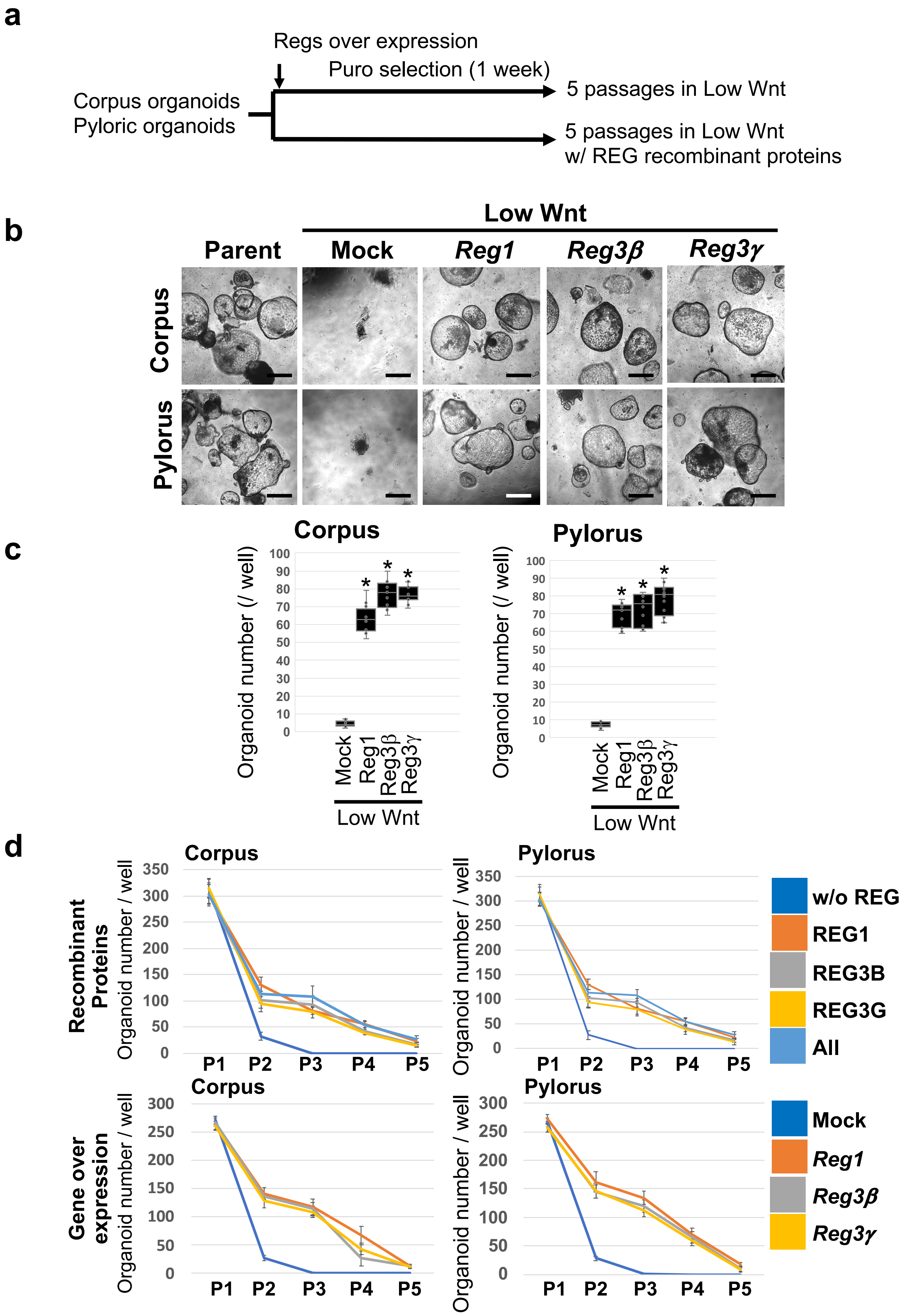


**Supplementary Figure 9.** (a) Experimental outline to confirm the contribution of *Reg* family genes to the regeneration of normal stomach epithelia. (b) Representative images of normal stomach organoids overexpressing *Reg1*, *Reg3β* or *Reg3γ* grown under low Wnt conditions. Scale bar = 200 µm. (c) Organoid quantification under low Wnt conditions. * $p < 0.05$. (d) Organoid quantification during long-term passage (~ P5). At each passage, 1 well was split into 3 wells.

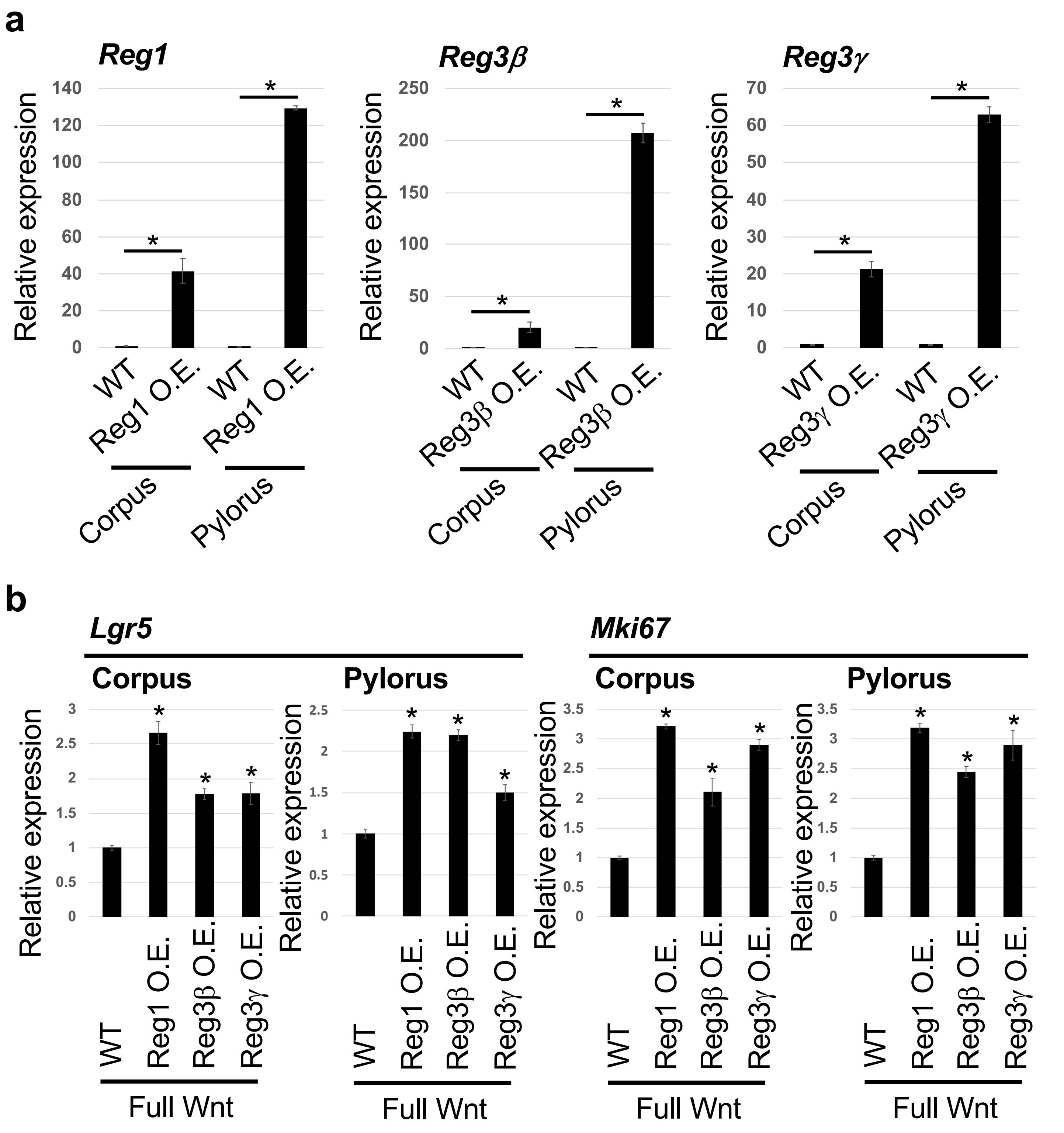


**Supplementary Figure 10.** (a) Confirmation of Reg gene expression levels by qPCR. * $p < 0.05$. (b) Expression levels of *Lgr5* and *Mki67* in Reg-overexpressing normal stomach organoids. * $p < 0.05$.

**Supplementary Table 1.**

**gRNA target sequences**

| gRNA name | Sequence |
|---|---|
| mApc_gRNA_MGLibA_04601 | AGATCCTTCCCGACTTCCGT |
| mApc_gRNA | CCGACTTCCGTAGGAGCGAA |
| mAlk_gRNA_MGLibA_03880 | TACACTTACCATATCGACTG |
| mAlk_gRNA | AGGAGAGCTCTGGTCGCCGA |
| mBclaf3_gRNA_MGLibA_02017 | CTCGTGTGAATTCGTATCAT |
| mBclaf3_gRNA | CGGGAGCTGCGCGACGGGCG |
| mPrkra_gRNA_MGLibA_42784 | TACTCGTGCAATACCTGAAT |
| mPrkra_gRNA | TGGCCGGCTCTTCCGCGCCG |
| hAPC_gRNA_1 | TTAGGGTTCAACTACACGAA |
| hAPC_gRNA_2 | CGAAGGCTGACAAGTCATCT |
| hALK_gRNA_1 | GAACATTGTTCGCTGCATTG |
| hALK_gRNA_2 | TGCAGCGAACAATGTTCTGG |
| hBCLAF3_gRNA_1 | ATCCAAAAAGACCCGTCGCG |
| hBCLAF3_gRNA_2 | AAGACCCGTCGCGTGGAGAA |
| hPRKRA_gRNA_1 | GGCGAAACATAGAGCTGCAG |
| hPRKRA_gRNA_2 | TATTCGTGTAATACCTGAAT |

**QPCR Primer list**

| Gene name | Forward | Reverse |
|---|---|---|
| *GapDH* | GGTGAAGGTCGGTGTGAACG | CTCGCTCCTGGAAGATGGTG |
| *Lgr5* | CAGTGTTGTGCATTTGGGGG | CAAGGTCCCGCTCATCTTGA |
| *Axin2* | AGGAGCAGCTCAGCAAAAAG | GCTCAGTCGATCCTCTCCAC |
| *Alk* | ACTGACATCCTCGCTTCTGAA | ATACGTTTCCTCTCAAAACCCC |
| *Bclaf3* | TCCACAAAGTTAAAGCAGATAGTTT | TCGCCTGTGAAATTCTGGATCT |
| *Prkra* | ACAGCGGGACCTTCAGTTTG | TCATGCCGTACTCGTGCAAT |
| *Reg1* | GCTGAAGAAGACCTGCCATC | AGATCTGCATCAGCCCAAGT |
| *Reg3* $\beta$ | CTGCCTTAGACCGTGCTTTC | AGGGCAACTTCACCTCACAT |
| *Reg3* $\gamma$ | AACAGAGGTGGATGGGAGTG | GTGATTGCCTGAGGAAGAGG |
| *Mki67* | AGCACAAAGAGACGGTCTAAGA | CTCTGCCTCGTGACTGTGTT |
| *Gif* | AAGCACAGCGCAAAAACTCC | GCAACCCCTTCATCCAAAGG |
| *Muc5ac* | GTGGTTTGACACTGACTTCCC | CTCCTCTCGGTGACAGAGTCT |
| *Cd44* | ACCTTGGCCACCACTCCTAAT | CCCTTCTGTCACATGGGAGTC |
| *Ctnnb1* | CGTGCAATTCCTGAGCTGACA | GCGTACAATGGCAGACACCATC |
